# Identification and Regulation of a Hepatic Lipogenic Metabolon

**DOI:** 10.64898/2025.12.30.696908

**Authors:** Xiaotong Zhu, William B. McKean, Tam Tran, Xiaorong Fu, Jay D. Horton, Chai-Wan Kim

## Abstract

De novo lipogenesis (DNL) plays a key role in the excessive fat accumulation present in Metabolic Dysfunction-Associated Steatotic Liver Disease (MASLD). Most mechanistic studies and experimental strategies for improving hepatic steatosis in MASLD have focused on the transcriptional regulation of enzymes involved in DNL and triglyceride (TG) synthesis. Here, we provide evidence for a post-translational mechanism that enhances fatty acid (FA) and TG synthesis through the assembly of a multi-protein lipogenic metabolon in liver. Under anabolic conditions, acetyl-CoA carboxylase 1 (ACC1) interacts with additional key enzymes in the DNL and TG synthesis pathway. Immunofluorescence and electron microscopy reveal that this lipogenic metabolon localizes around lipid droplets (LDs) and in proximity to mitochondria and LD interfaces in the anabolic state. The formation of the lipogenic metabolon facilitates the efficient transfer of FA synthesis intermediates to enhance lipogenic flux. These findings uncover a new nutrient-responsive, post-translational regulatory mechanism for hepatic lipogenesis and highlight the lipogenic metabolon as a potential therapeutic target for metabolic liver diseases.

## INTRODUCTION

*De novo* lipogenesis (DNL) is the process in which fatty acids (FAs) are synthesized in cells from acetyl-CoA, primarily derived from carbohydrate metabolism ^1^. Key transcriptional regulators of DNL, including SREBP-1c and ChREBP, are tightly regulated by hormones such as insulin ^2,3^ and glucagon ^2^, and through post-translational modifications of the DNL enzymes including phosphorylation ^2^ and acetylation ^4,5^. In a healthy liver, DNL contributes to less than 5% of total hepatic triglycerides (TGs) ^6–8^. However, in Metabolic Dysfunction-Associated Steatotic Liver Disease (MASLD), this contribution rises to 25-30% ^9–11^, due to insulin resistance, which leads to an increase in hepatic FA and TG synthesis ^2,12–14^. Excessive TG accumulation in liver is associated with progression to metabolic-associated steatohepatitis (MASH), cirrhosis, and hepatocellular carcinoma ^15–17^. With over one-third of U.S. adults affected by MASLD ^18^, it is crucial to elucidate the mechanisms driving aberrant hepatic FA synthesis in insulin-resistant states to identify potential therapeutic targets for FA-driven metabolic disorders.

Our previous studies have shown that the excessive activation of SREBP-1c initiates the transcription of genes encoding the enzymes necessary for FA synthesis, as well as the enzyme catalyzing the first step in glycerolipid synthesis ^12,13^. This transcriptional program is a major driver of elevated FA and TG synthesis in insulin-resistant states ^2,12,13^, subsequently leading to hepatic steatosis, increased VLDL secretion, and hypertriglyceridemia ^13,19–24^. The requirement of SREBP activation for the development of hepatic steatosis and hypertriglyceridemia was demonstrated by studies in which SCAP, a protein required for SREBP activation, was deleted in obese, insulin resistant mice (*ob/ob* mice) that normally have very high levels of hepatic FA and TG synthesis. SCAP deletion completely normalized rates of FA synthesis, liver TGs, VLDL secretion, and plasma TG levels ^24^ in *ob/ob* mice and hamsters. These results demonstrate that enhanced DNL is a major driver of steatosis in insulin-resistant livers.

FA and TG synthesis requires sequential enzymatic reactions distributed across multiple subcellular compartments. ATP citrate lyase (ACL) produces acetyl-CoA from citrate, acetyl-CoA synthetase 2 (ACSS2) synthesizes acetyl-CoA from acetate, acetyl-CoA carboxylases (ACC) 1 and 2 convert acetyl-CoA to malonyl-CoA, and fatty acid synthase (FAS) uses malonyl-CoA and acetyl-CoA to generate palmitate (C16:0). These enzymes (except for ACC2 that is predominantly localized to the outer mitochondrial membrane) are predominantly localized in the cytosol ^25^. Palmitate can then be further desaturated by stearoyl-CoA desaturase (SCD1) or elongated by ELOVL fatty acid elongase 6 (ELOVL6) before further desaturation, but unlike the enzymes involved in the synthesis of palmitate, these proteins reside in the endoplasmic reticulum (ER) ^26,27^. Additionally, the first enzyme in glycerolipid synthesis, glycerol-3-phosphate acyltransferase 1 (GPAM), which acylates glycerol-3-phosphate to initiate TG and phospholipid synthesis is located in the mitochondria ^28^. How the FA and glycerolipid synthesis intermediates are efficiently transferred among these spatially distributed enzymes to sustain continuous and efficient lipid synthesis remains unclear.

One conceptual framework that could address this challenge is the metabolon hypothesis. In 1970, A.M. Kuzin first introduced the concept of physical enzyme–enzyme complexes. This idea gained momentum in 1972 when P.A. Srere applied it to enzymes of the citric acid cycle, highlighting their potential functional interactions ^29^. Srere further expanded on this idea by coining the term “metabolon” during a 1985 lecture in Debrecen, Hungary. He defined it as “a supramolecular complex of sequential metabolic enzymes and cellular structural elements” ^30^, emphasizing the functional coordination within cellular metabolic pathways. This concept has been applied to explain the spatial organization of metabolic pathways such as the citric acid cycle ^31,32^ and glycolysis ^33^, prompting us to examine whether hepatic FA and TG synthesis might also be coordinated through spatial organization of lipogenic enzymes.

Here, we show that under anabolic conditions—such as high-carbohydrate feeding and insulin resistance—FA and TG synthesis enzymes assemble into a functional lipogenic metabolon. This complex enhances lipogenic flux by coordinating the sequential enzymatic reactions required to synthesize the end product. This metabolon, including accessory proteins MIG12 and SPOT14 (S14) ^34^, predominantly localizes around lipid droplets (LDs) in primary hepatocytes. Together, these findings reveal a dynamic multi-enzyme complex that facilitates FA and TG synthesis and provides new insights into the spatial organization and coordinated regulation of hepatic lipid synthesis.

## RESULTS

### Identification of the lipogenic complex in liver

The first indication that a lipogenic complex might exist grew out of investigations focused on determining the function of a poorly characterized protein, S14. S14 is highly expressed in lipogenic tissues ^35^, and its gene transcription is regulated by SREBP-1c ^36,37^, carbohydrate-responsive element-binding protein (ChREBP) ^38^, and thyroid hormone receptor (TR) ^39^, which suggests that it may play a role in lipogenesis. Deletion of S14 in mice reduced lipid synthesis in liver ^34^ and lactating mammary glands ^40^, although its exact function in lipogenesis has not been elucidated. To further define the function of S14, we used an adenovirus that expressed S14 with a Flag tag in mice and carried out immunoprecipitation of S14 from liver lysates to identify potential interacting proteins. LC/MS analysis revealed that MIG12, ACC1, FAS, and β-tubulin all co-immunoprecipitated with S14 (Figure 1A). MIG12, a protein we previously showed that promotes the polymerization of ACC1 and ACC2 ^41^, shares 32% sequence identity with S14 ^41^ and can form a heterodimer with S14 ^34^. To further confirm that the association of these enzymes co-immunoprecipitated with S14 reflects physical interaction in cells, we performed immunofluorescence staining in primary hepatocytes isolated from rats refed a fat-free, high-carbohydrate diet (FFD). ACC1, MIG12, and S14 prominently co-localized around LD–associated ER regions. Quantitative analysis using Pearson’s correlation (above Costes threshold) demonstrated strong spatial overlap among the three enzyme pairs, with ACC1–S14 (R = 0.73), ACC1–MIG12 (R = 0.52), and S14–MIG12 (R = 0.52) (Supplemental Table 1). These data indicate that ACC1, S14, and MIG12 occupy shared subcellular domains under conditions of high lipogenesis, supporting the existence of a coordinated lipogenic enzyme complex in hepatocytes (Figure 1B). Representative ROIs and pixel-intensity histograms are shown in Figure S1, and detailed colocalization metrics, including Manders’ coefficients and Costes statistics, are provided in Supplemental Table 1.

**Figure 1.**
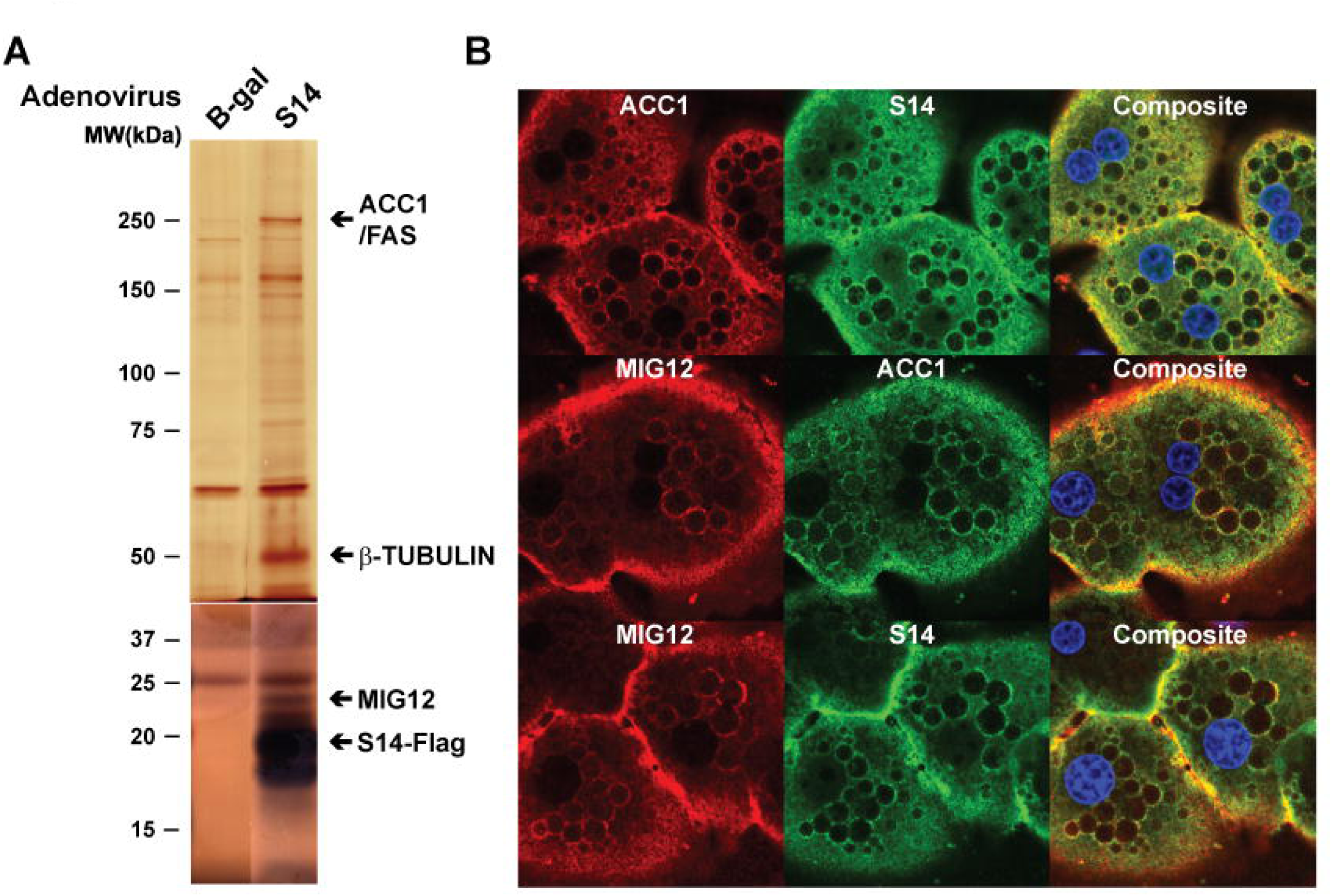
Identification of the lipogenic complex in liver. (A) An adenovirus expressing S14-Flag was injected into mice, and liver lysates were prepared 3 days later. Immunoprecipitation was performed using anti-Flag antibodies. Eluates from the S14-Flag IP and an identically processed β-gal–injected control were separated on 8% and 15% SDS–PAGE gels, followed by silver staining. Protein bands that appeared specifically in the S14-Flag lanes were excised and analyzed by LC–MS/MS to identify potential S14-associated proteins. (B) Primary hepatocytes were prepared from rats refed a FFD and plated on collagen-coated coverslips. Immunofluorescence confocal imaging was performed as described in Methods. Co-localization of ACC1 with MIG12 and S14 was confirmed by confocal microscopy using anti-mouse monoclonal ACC1, anti-MIG12 rabbit polyclonal antibodies, and anti-S14 rabbit polyclonal antibodies. Quantitative co-localization analysis was performed using Pearson’s correlation (above Costes threshold), and full analytical metrics, including Manders’ coefficients and Costes significance testing, are provided in Figure S1 and Supplemental Table 1.

### Co-localization of proteins involved in lipogenesis around LDs in hepatocytes

To investigate potential additional interactions among lipogenic proteins, we used antibodies recognizing various lipogenic enzymes in primary hepatocytes to assess co-localization by confocal immunofluorescence. As shown in Figure 1, co-localization was quantified using Pearson’s correlation coefficient (Pearson’s R) in LD-enriched regions where the enzyme signals were concentrated. Immunofluorescence imaging of primary hepatocytes isolated from rat livers refed a FFD demonstrated that ACC1 co-localized with ACL (R ≈ 0.83), ACSS2 (R ≈ 0.46), and FAS (R ≈ 0.55) around LDs, despite their known localization in the cytosol. GPAM, a mitochondrial outer membrane protein involved in glycerolipid synthesis, also co-localized with ACC1 in LD-associated regions (R ≈ 0.51). In contrast, the gluconeogenic enzyme PEPCK, which is unrelated to lipogenesis, showed almost no overlap with ACC1 (R ≈ 0.11), likely because it remains confined to the cytosol and does not redistribute to LD-proximal regions (Figure 2A, Supplemental Table 2). Comprehensive testing metrics, including Manders’ coefficients and Costes significance testing, are provided in Supplemental Table 2 and further support the robust spatial association of ACC1 with multiple cytosolic lipogenic enzymes, whereas the non-lipogenic negative control, PEPCK, showed only minimal overlap with ACC1. Adipose differentiation-related protein (ADRP), an LD-resident protein, marked LDs where ACL, ACC1, and FAS also localized in primary hepatocytes from refed rats (Figure 2B). Three-dimensional imaging further confirmed the localization of ACC1 surrounding LDs in primary hepatocytes from refed rats (Figure 2C). Additional imaging across various fasting and refeeding timepoints suggested that ACC1-containing complexes assemble progressively during metabolic activation and accumulate around LDs in rat primary hepatocytes (Figure S2).

**Figure 2.**
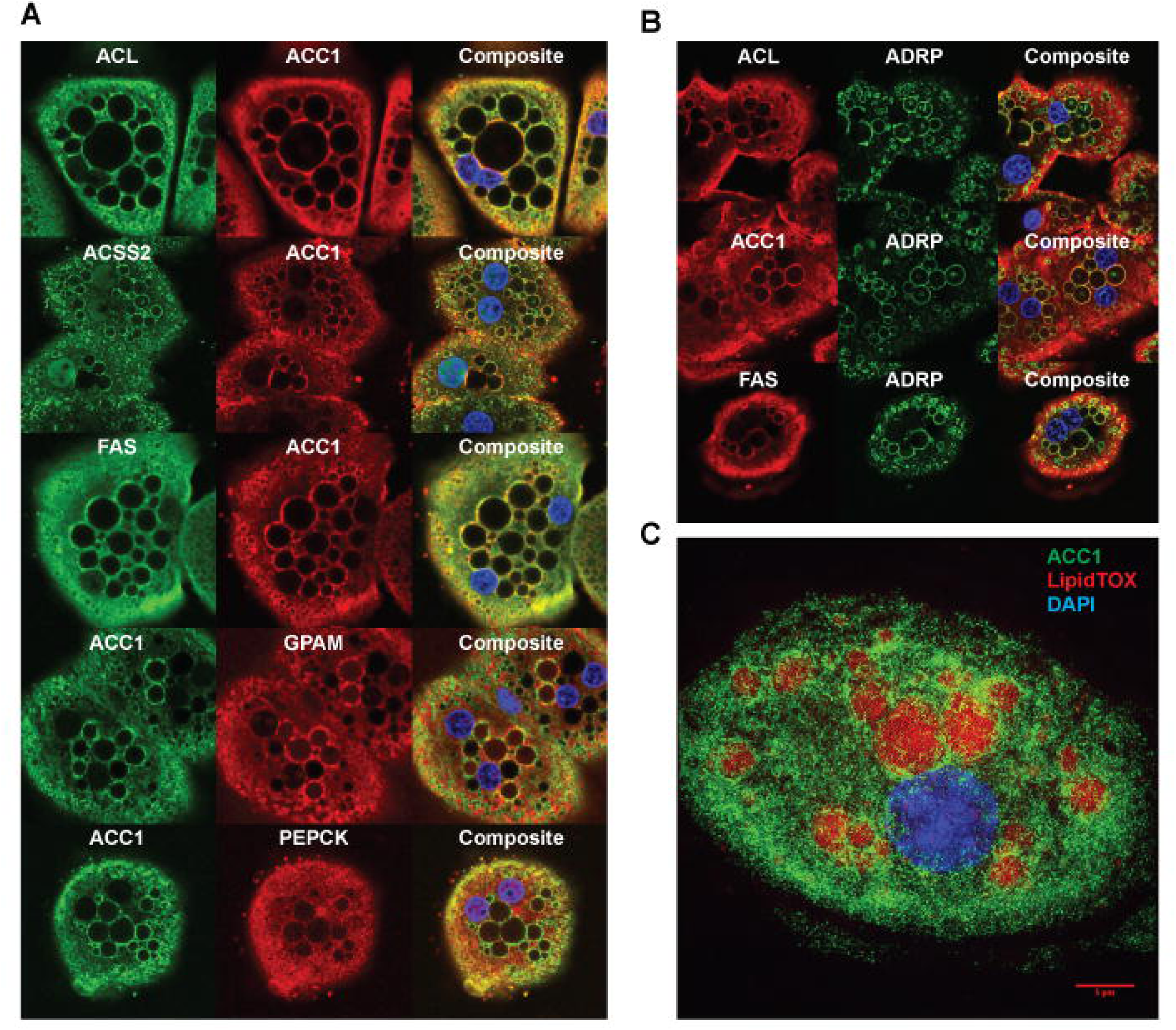
Co-localization of proteins involved in lipogenesis around LDs in hepatocytes. (A) Co-localization of enzymes involved in lipogenesis in primary hepatocytes from rats refed a FFD using confocal microscopy as described in Methods. Quantitative co-localization analysis was performed using Pearson’s correlation (above Costes threshold), and full analytical metrics are provided in Supplemental Table 2. (B) Localization of enzymes involved in lipogenesis around LDs in primary hepatocytes from rats refed a FFD using confocal microscopy as described in Methods. Co-localization analysis was performed using Pearson’s correlation as in panel A, with results summarized in Supplemental Table 2. (C) A three-dimensional image stack was acquired using an OMX microscope (z-step size: 120 nm) to visualize the indicated proteins around lipid droplets, as described in Methods.

### Co-localization of proteins involved in lipogenesis at the interface of mitochondria and LDs in hepatocytes

Electron microscopy (EM) with immunogold labeling using antibodies recognizing ACC1 or FAS was used to determine their subcellular localization in primary hepatocytes from refed rats. Consistent with the immunofluorescence images (Figure 2C; Figure S2), EM revealed that ACC1 localized predominantly around LDs (Figure 3A). This finding was further supported by immunofluorescence imaging and EM performed on the same hepatocyte (Figure S3A). Double immunogold labeling using different sizes of gold particles for ACC1 and FAS demonstrated their co-localization around LDs (Figure 3B; Figure S3B–C). Although ACC1 and FAS were occasionally observed near the ER, they were much more consistently enriched around LDs (Figure S3D). This pattern aligned with staining of the ER marker protein disulfide isomerase (PDI), which showed only limited overlap with ACC1, in contrast to the strong co-localization observed between ACC1 and LDs (Figure S3E).

**Figure 3.**
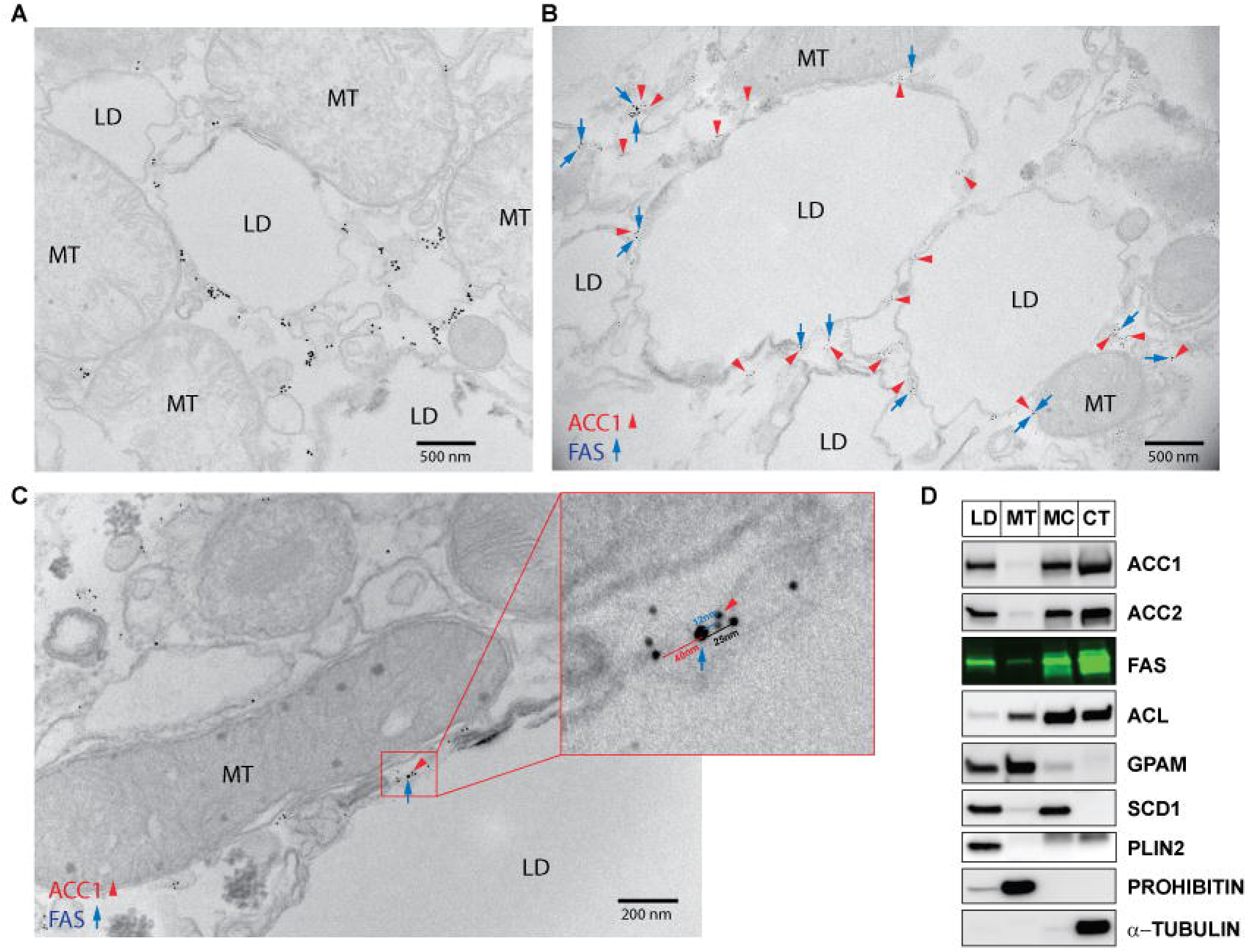
Co-localization of proteins involved in lipogenesis at the interface of mitochondria and LDs in hepatocytes. (A) Primary hepatocytes were prepared from rats refed a FFD and localization of ACC1 was determined using immunogold-EM. The 1.4 nm nanogold particles were enlarged by gold enhancement for clear visibility. Detailed protocol is described in Methods. (B) Co-localization of ACC1 and FAS around LDs in primary hepatocytes from rats refed a FFD using immunogold-EM. Size of gold particles: ACC1 - 5 nm; FAS - 10 nm. (C) Co-localization of ACC1 and FAS between mitochondria and LDs in primary hepatocytes from rats refed the FFD using immunogold-EM. Size of gold particles: ACC1 - 5 nm; FAS - 10 nm. (D) Subcellular fractionation was performed with liver lysates from rats refed the FFD to isolate LDs as well as mitochondria, microsomes, and cytoplasm. Detailed protocols for the subcellular fractionation are described in Methods. Proteins from each fraction were subjected to SDS-PAGE and immunoblot analysis with the indicated antibodies. PLIN2, prohibitin, and α-tubulin were used as subcellular localization markers for lipid droplets, mitochondria, and cytosolic proteins, respectively.

Notably, interactions between ACC1 and FAS were also observed at interfaces between mitochondria and LDs (Figure 3C). Subcellular fractionation of liver lysates from FFD-refed rats further demonstrated that ACL, ACC1, ACC2, FAS, SCD1, and GPAM were present in both LD and mitochondrial fractions, regardless of their canonical localizations in the cytoplasm, mitochondria, or ER (Figure 3D). EM frequently revealed mitochondria positioned adjacent to LDs (Figure S3F), and GPAM localization was subsequently confirmed to be on the outer mitochondrial membrane, including sites where mitochondria interface with LDs (Figure S3G). Subcellular fractionation of liver lysates from FFD-refed rats further confirmed this finding, whereas mitochondrial inner membrane proteins, such as prohibitin and cytochrome c, were found only in the mitochondrial fraction (Figure S3H). Combined, these findings demonstrate that lipogenic proteins co-localize around LDs and at mitochondria–LD contact sites under high lipogenesis conditions.

### Active dephosphorylated ACC promotes polymerization, enhances lipogenesis, and increases LD localization

ACC activity is acutely regulated through phosphorylation and dephosphorylation. AMP-activated protein kinase (AMPK) phosphorylates ACC at Ser-79 ^42,43^, inhibiting its enzymatic activity through largely undefined mechanisms ^44,45^. Fasting or glucagon treatment increases AMPK activity, whereas insulin suppresses AMPK activity and stimulates the Mg²⁺-dependent protein phosphatase 2A (PP2A), which dephosphorylates ACC, thereby increasing its activity ^46–50^. During refeeding, ACC is converted from its phosphorylated inactive form to its dephosphorylated, active form due to stimulation of protein phosphatase activity. This dephosphorylation activates ACC and DNL in hepatocytes, allowing excess glucose to be converted into FAs and TGs for storage^51,52^.

We next determined whether the phosphorylated state of ACC affected the formation of the lipogenic complex. For this study, we treated rat primary hepatocytes with 5-aminoimidazole-4-carboxamide ribonucleoside (AICAR), an AMPK agonist, to induce ACC phosphorylation ^53^. The primary hepatocytes were incubated with 1 mM AICAR and cell lysates were prepared at 0, 1.5, and 3 hours after treatment. SDS-PAGE immunoblot analysis of cell lysates showed an increase in phosphorylated ACC levels in the AICAR-treated cells compared to the untreated cells; however, the amount of total ACC protein was unchanged (Figure 4A). Blue native PAGE analysis of cell lysates demonstrated that the polymerized, higher molecular weight form of ACC1 was present only in untreated primary hepatocytes and was disrupted in AICAR-treated hepatocytes (Figure 4A). Even with MIG12 overexpression, which robustly promotes ACC1 polymerization, AICAR reduced the higher molecular weight form of ACC1 to baseline (Figure 4B). Consistent with reduced polymerization, FA synthesis measured by ³H-acetate incorporation into newly synthesized FAs fell ∼6-fold within 30 minutes of AICAR treatment (Figure 4C), while FA-synthesis enzyme levels remained unchanged (Figure S4A). Collectively, these results imply that the rapid decrease in DNL results from AMPK-mediated ACC phosphorylation, which disrupts ACC1 polymerization.

**Figure 4.**
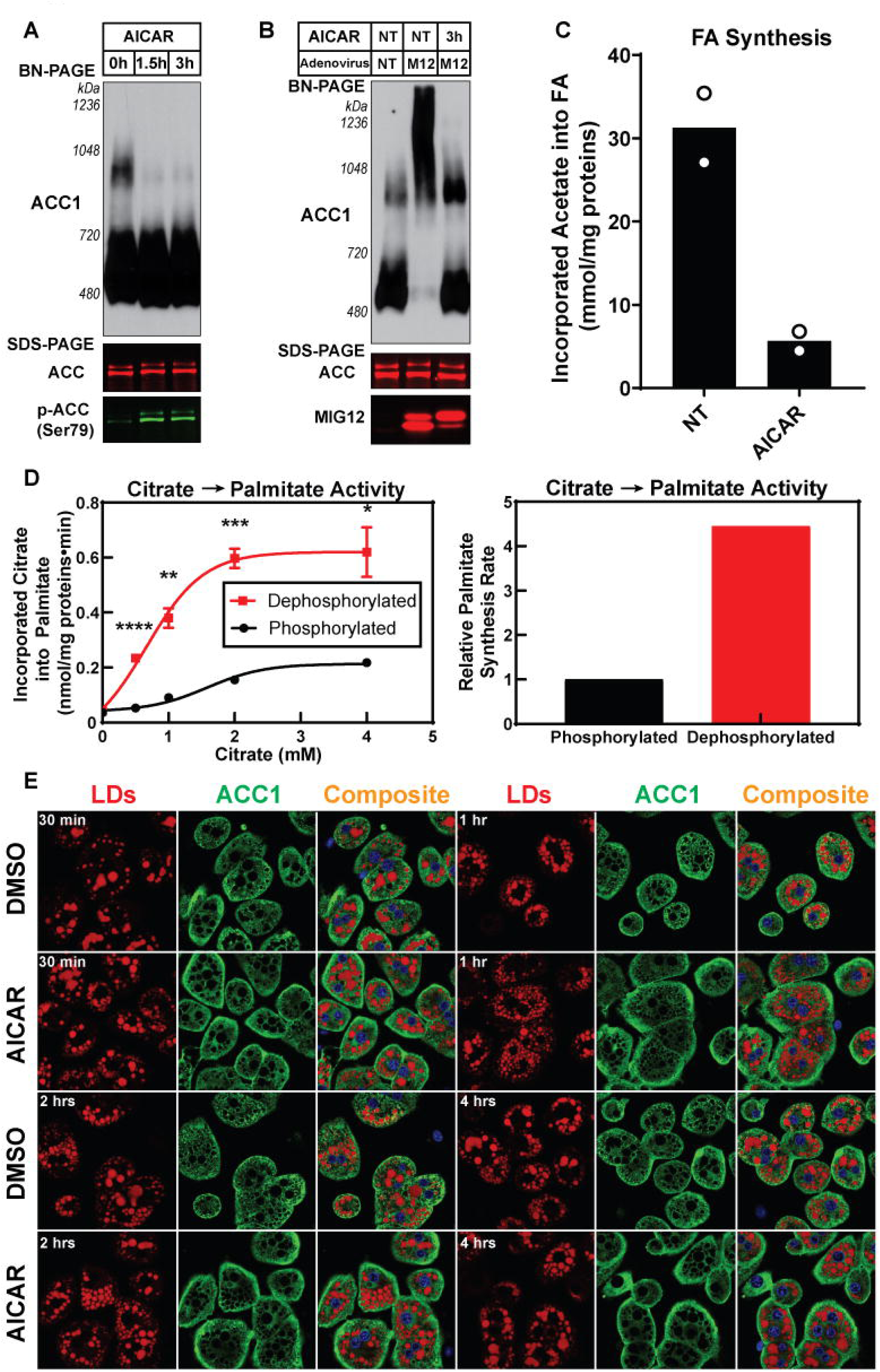
Active dephosphorylated ACC promotes polymerization, enhances lipogenesis, and increases LD localization. (A) Primary hepatocytes from rats refed a FFD were treated with 1 mM AICAR for 0, 1.5, and 3 hours. Equal aliquots of cell lysate proteins were loaded and analyzed by BN-PAGE (above) or SDS-PAGE (bottom) for immunoblot analysis. An anti-ACC1 antibody was used to perform the immunoblot analysis of BN-PAGE. For SDS-PAGE, ACC immunoblots were performed using a fluorescently labeled streptavidin conjugate. (B) Primary hepatocytes from rats refed a FFD were isolated. One set of primary hepatocytes were untreated, while the other two sets were infected with an adenovirus expressing MIG12. After 12 hours, 1 mM AICAR was added, and cells were harvested 3 hours later. Cell lysates were prepared and subjected to BN-PAGE and SDS-PAGE for immunoblot analysis. An anti-ACC1 antibody was used to perform the immunoblot analysis of BN-PAGE. For SDS-PAGE, a fluorescently labeled streptavidin conjugate to detect ACC and an anti-MIG12 antibody were used for immunoblot analysis. (C) Primary rat hepatocytes prepared as in (B) were incubated with 1 mM AICAR for 30 minutes and FA synthesis rates were measured *ex vivo* using ^3^H-acetate as described in Methods. Individual data points are shown; bars indicate the mean (n = 2 technical replicates from one hepatocyte preparation). (D) Liver cytosol from rats refed a FFD were pre-incubated with 1 mM MnCl₂ in the presence or absence of a phosphatase inhibitor cocktail for 15 min at 30°C, then incubated with the indicated concentrations of Na-citrate buffer (pH 7.1) for 15 min at 37°C. Palmitate synthesis (left) was measured using ¹⁴C-citrate. Palmitate synthesis rates (right) were calculated from the maximal velocity (Vmax) of the enzymatic reaction. Data represent mean ± SEM from n = 3 technical replicates. Statistics: multiple two-tailed unpaired Student’s t-tests with Holm–Šidák correction, comparing phosphatase inhibitor–treated (phosphorylated) vs untreated (dephosphorylated) conditions at each citrate concentration (*P < 0.05, **P < 0.01, ***P < 0.001, ****P < 0.0001). The detailed protocol for the enzymatic assay is described in Methods. (E) Rat primary hepatocytes were isolated from rats fed a FFD for 2 days and incubated in medium containing either (i) 25 mM glucose, 100 nM insulin, and 1% DMSO or (ii) 5 mM glucose and 1 mM AICAR. Following the indicated times, cells were stained with anti-ACC1 antibodies and LipidTOX as described in Methods.

To further explore the role of dephosphorylated ACC in promoting DNL *in vitro*, we blocked endogenous phosphatase activity in liver cytosol using a phosphatase inhibitor cocktail (Figure 4D). Immunoblot analysis confirmed that the inhibitor prevented ACC dephosphorylation, whereas λ-phosphatase enhanced dephosphorylation (Figure S4B). Compared with the inhibitor-treated group, untreated liver cytosol, which underwent dephosphorylation during incubation, showed a 4-fold increase in palmitate synthesis rates in a citrate concentration-dependent manner (Figure 4D). Furthermore, in primary hepatocytes from refed rats, ACC1 localized around LDs under DMSO control, consistent with Figures 1–3, but this LD-associated pattern was significantly reduced with AICAR treatment (Figure 4E).

Taken together, these findings demonstrate that the polymerization of ACC, mediated by dephosphorylation, is critical for its localization with other lipogenic enzymes around LDs and for regulating DNL in hepatocytes.

### The interaction of ACC1 with lipid synthesis-associated proteins is enhanced when ACC is in the activated state

As demonstrated above, ACC1 polymerizes and co-localizes with other lipogenic proteins around LDs under high DNL conditions. To determine how ACC1 interactions with other proteins change upon activation, we performed ACC1 immunoprecipitation from mouse liver lysates incubated with 5 mM MgCl₂ at 30°C for 30 minutes with or without a phosphatase inhibitor cocktail. With endogenous phosphatases active, ACC1 co-immunoprecipitated ACC2, ACSS2, FAS, ACL, GPAM, MIG12, S14, and SCD1 (Figure 5A). Notably, β-actin, a cytoskeletal protein, also showed significant binding to ACC1, suggesting that cytoskeletal components might play a role in the formation of the lipogenic complex. In contrast, these interactions were markedly reduced when endogenous phosphatases were inhibited, suggesting that ACC1 interacts more strongly with other proteins in a phosphatase-active state. Proteins unrelated to lipid synthesis, such as PEPCK, prohibitin, cytochrome c, and PDI, did not bind ACC1 under either condition.

**Figure 5.**
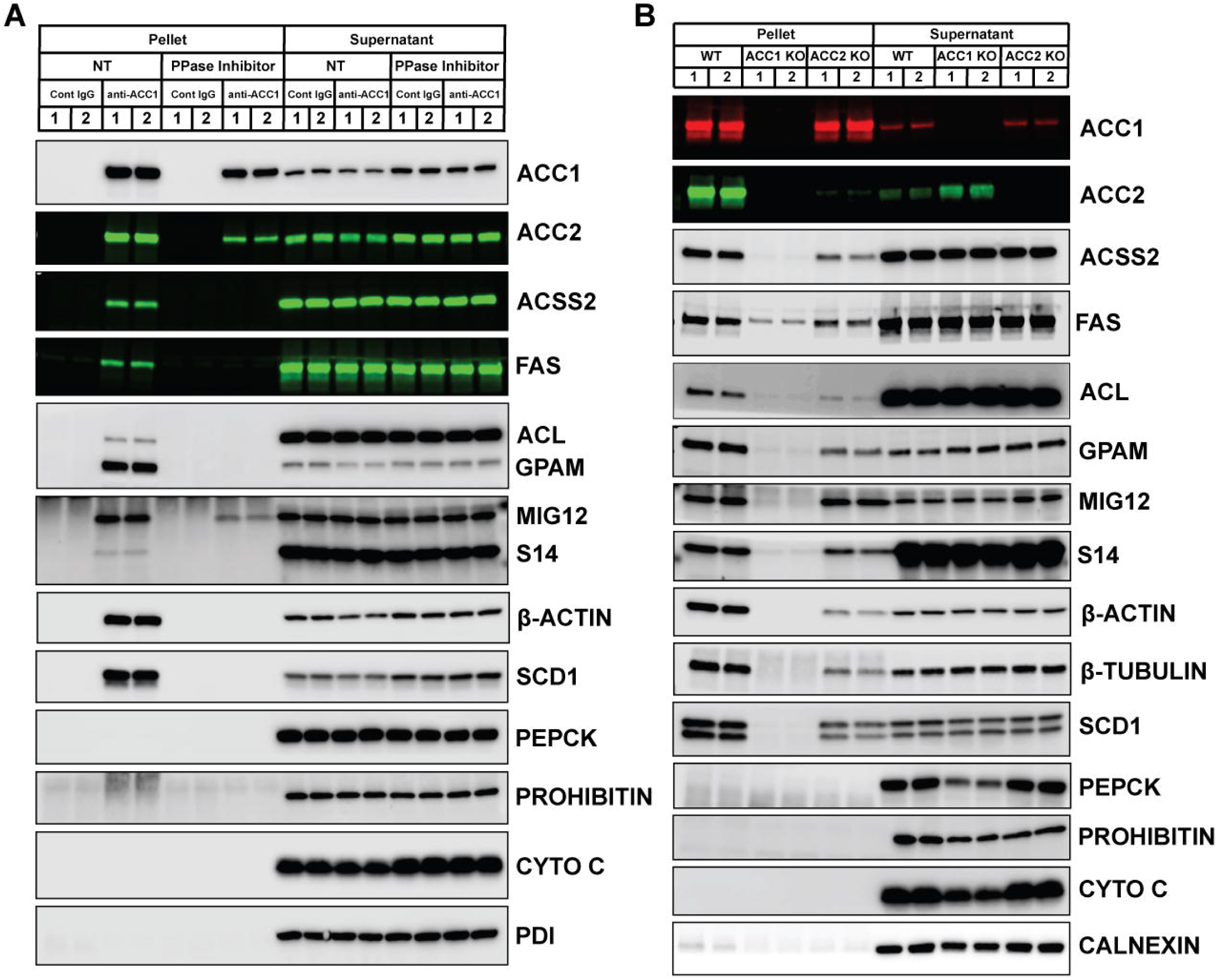
The interaction of ACC1 with lipid synthesis-associated proteins is enhanced when ACC is in the activated state. (A) Liver lysates from refed mice were treated with or without a phosphatase inhibitor cocktail. The ACC1 antibody was incubated with the lysate for 1 hour at RT and ACC1 was immunoprecipitated. IgG was used as the control. The immunoprecipitation protocol is described in Methods. Supernatant indicates the unbound fraction collected after bead pelleting; it was mixed with SDS sample buffer and immunoblotted alongside the bead-bound material. (B) ACC1 immunoprecipitation was carried out using lysates prepared from refed wild-type, ACC1 hepatocyte knockout, or ACC2 hepatocyte knockout mouse livers, respectively.

Notably, ACC2 maintained strong binding to ACC1 even in the presence of phosphatase inhibitors, suggesting that ACC1 and ACC2 form a relatively stable heteromer. These observations are supported by immunoprecipitation of liver lysates from hepatocyte-specific ACC1 or ACC2 knockout mice, which revealed a clear reciprocal interaction between the two enzymes (Figure S5A). Blue native PAGE analysis further supported this model: whereas ACC1 migrated similarly in wild-type and ACC2-deficient livers, ACC2 showed a markedly altered pattern in ACC1-deficient livers. In normal liver, ACC2 predominantly appeared as a dimer or tetramer, but in the absence of ACC1, ACC2 shifted toward higher–molecular-weight, more polymerized species (Figure S5B). Considering that hepatic ACC2 expression is substantially lower than ACC1, these findings suggest that, under physiological conditions, most endogenous ACC2 may exist as a heterodimer with ACC1.

Immunofluorescence analysis in cultured cells provided additional evidence for the ACC1–ACC2 heterodimer. ACC1 was detected predominantly in the cytosol when expressed alone (Figure S5C). However, when co-transfected with ACC2, a substantial fraction of ACC1 colocalized with ACC2 at the mitochondria (Figure S5D). In addition to this mitochondrial pool, ACC1 also displayed cytosolic localization that did not overlap with ACC2. These observations indicate that ACC1 is present in the cytosol but also at the mitochondrial membrane as a complex with ACC2.

To determine whether endogenous ACC2 influences ACC1-mediated protein interactions, we immunoprecipitated ACC1 from refed wild-type and ACC2 hepatocyte-specific knockout livers, using ACC1-deficient mice as a negative control. Co-precipitation of ACL, ACSS2, FAS, S14, SCD1, GPAM, β-actin, and β-tubulin was reduced to varying degrees in ACC2-deficient livers compared with wild-type (Figure 5B), suggesting that ACC2 contributes to the stability or assembly of ACC1-associated protein complexes. MIG12 binding to ACC1 was unaffected by phosphatase inhibition (Figure 5A) or by the absence of ACC2 (Figure 5B), consistent with its direct association with ACC1. As expected, PEPCK, prohibitin, cytochrome c, and calnexin were not co-immunoprecipitated with ACC1 in any group.

Given that ACL, ACSS2, ACC, FAS, SCD1, and the mitochondrial GPAM participate in ACC1-associated complexes, we next measured TG synthesis under phosphatase active versus phosphatase-inactive conditions. Glycerol 3-phosphate (G3P) was used as a substrate with excess oleoyl-CoA added to liver homogenates from refed rats to test whether TG synthesis via GPAM is enhanced when endogenous phosphatases are active, a condition that favors lipogenic complex formation (Figure S5E). The results demonstrated that TG synthesis was significantly higher under phosphatase-active conditions and increased further with rising MgCl₂ concentrations, paralleling MgCl₂-dependent ACC1 dephosphorylation (Figure S4B), whereas under phosphatase inactive conditions, TG synthesis remained unchanged with MgCl₂ (Figure S5E).

These results demonstrate that phosphorylation–dephosphorylation regulates both FA and TG synthesis in part by controlling ACC1-mediated lipogenic complex formation.

### Lipogenic metabolon formation facilitates FA synthesis

To confirm the interactions between ACC1 and the other FA synthesis proteins, and to assess the regulatory effects of phosphorylation and dephosphorylation on the activities of individual FA synthesis enzymes *in vitro*, we conducted a comprehensive purification of recombinant ACSS2, ACC1, FAS, MIG12, and S14 in mammalian cells using Ni-NTA resin and gel-filtration chromatography (Figure 6A, S6A), and rPEPCK was purified as a negative control (Figure S6B). The recombinant protein mixture was dephosphorylated with λ-phosphatase and immunoprecipitation was performed with anti-ACC1 antibodies. Immunoblot analysis (Figure 6B) revealed that all other recombinant FA synthesis proteins co-immunoprecipitated with rACC1, whereas rPEPCK did not. These results confirm that recombinant ACC1 interacts with core lipogenic factors *in vitro* using recombinant proteins (Figure 6B).

**Figure 6.**
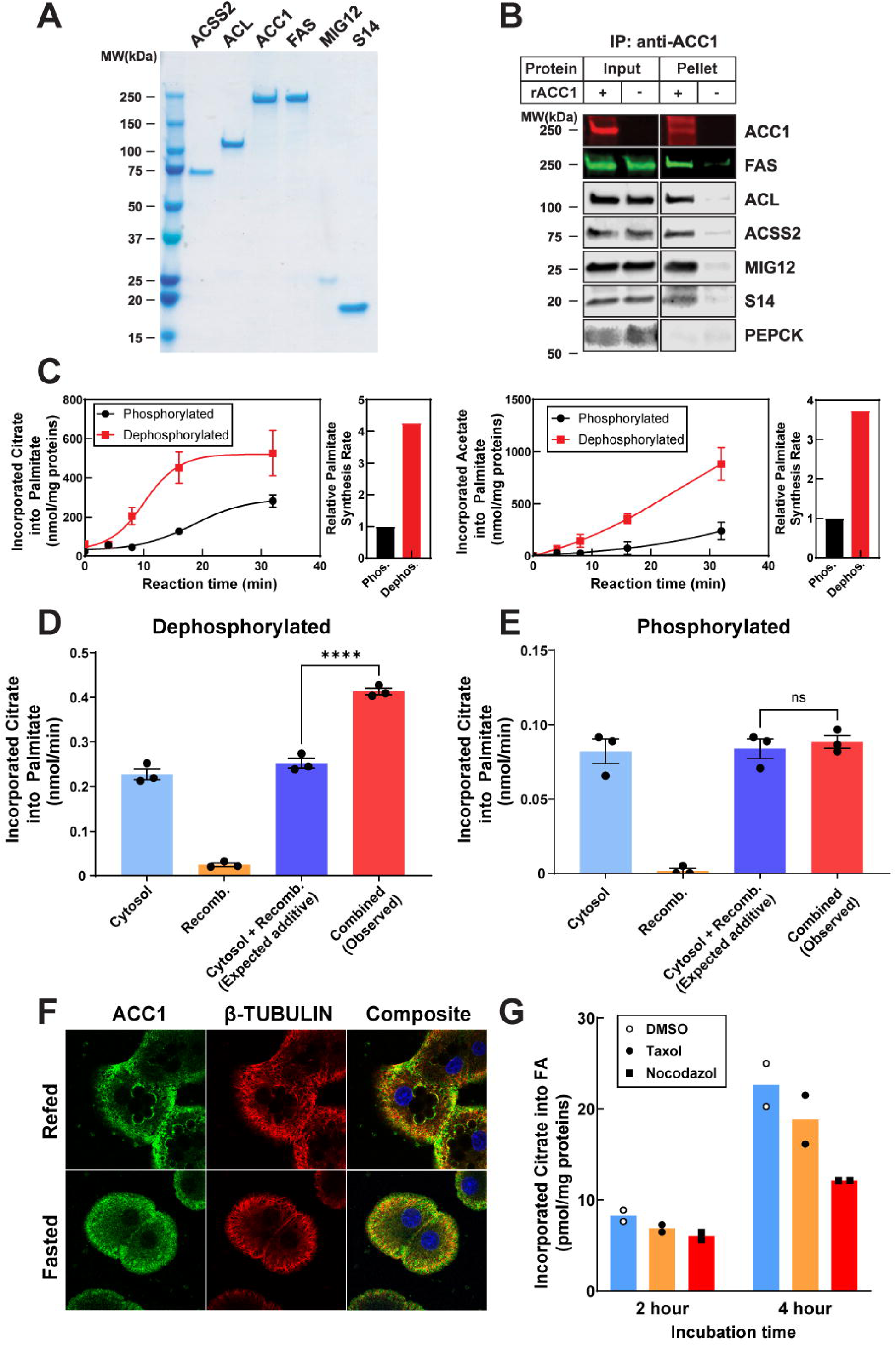
Lipogenic metabolon formation facilitates FA synthesis. (A) Human C-terminal His-Flag-tagged ACSS2, ACL, ACC1, FAS, MIG12, and S14 were individually overexpressed in mammalian HEK 293S GnTI^−^ cells and purified using Ni-NTA and gel filtration. Epitope tags were removed by TEV cleavage. Detailed protocols for purification were described in Methods. The final purified recombinant proteins were subjected to SDS-PAGE for Coomassie blue staining. (B) Recombinant protein mixtures were incubated with λ-phosphatase and 1 mM MnCl_2_ for 30 minutes at 30°C and then the protein mixture was incubated with 0.5 mM Na-citrate buffer for 20 minutes at 30°C. rACC1 was immunoprecipitated from the protein mixture using an anti-ACC1 antibody. Recombinant protein mixtures without rACC1 were used for control immunoprecipitations. Proteins were subjected to SDS-PAGE for immunoblot analysis. ACC1 immunoblots were performed using a fluorescently labeled streptavidin conjugate. (C) Enzymatic activity of reconstituted recombinant lipogenic enzyme modules was measured as described in Methods. The left panel shows FA synthesis activity measured from citrate using rACL–rACC1–rFAS, whereas the right panel shows FA synthesis activity measured from acetate using rACSS2–rACC1–rFAS. Reactions were performed under phosphorylated or dephosphorylated conditions as indicated. Data represent mean ± SEM from n = 3 technical replicates. Relative synthesis rates were calculated based on Vmax values. (D–E) Under dephosphorylated (D) or phosphorylated (E) conditions, ACL–ACC1–FAS activity was measured in mixtures of liver cytosol (from FFD-refed rats) and recombinant proteins (rACL, rACC1, and rFAS) and compared with the expected additive activity calculated from assays performed in cytosol alone and recombinant proteins alone in parallel. Data represent mean ± SEM from n = 3 technical replicates. Statistical analysis was performed using two-way ANOVA, revealing a significant interaction between phosphorylation state and mixing condition. Post hoc comparisons indicate that observed activity significantly exceeded the expected additive activity only under dephosphorylated conditions. *P < 0.05, **P < 0.01, ***P < 0.001, ****P < 0.0001. (F) Primary hepatocytes isolated from rats under refed or fasted conditions were fixed and stained for ACC1 and β-tubulin, followed by confocal microscopy as described in Methods. Images show representative localization patterns under each condition. (G) Primary hepatocytes from FFD-fed rats were cultured for 8 hours and then treated with DMSO, nocodazole, or taxol (20 μM) as indicated. Cells were subsequently incubated with ^14^C-labeled citrate for 2 or 4 hours, and citrate-derived FA synthesis was quantified as described in Methods. Data points represent individual measurements from duplicate experiments.

Next, we pre-incubated the individual recombinant proteins or their mixtures with λ-phosphatase to simulate the DNL activated environment as described above. Parallel groups without λ-phosphatase treatment were used as phosphorylated controls. Figure S6C shows that each purified protein exhibited functional activity, and interestingly, all tested lipogenic proteins displayed increased activity to varying degrees when dephosphorylated compared to their phosphorylated forms. However, rACC1 exhibited the greatest increase in enzymatic activity upon λ-phosphatase treatment compared with the other lipogenic enzymes. Next, we combined purified lipogenic enzymes, rACL-rACC1-rFAS or rACSS2-rACC1-rFAS, and measured FA synthesis from citrate or acetate, respectively. As predicted, rACL-rACC1-rFAS activity was nearly four times higher with λ-phosphatase treatment compared with untreated conditions (Figure 6C). Similarly, rACSS2-rACC1-rFAS activity increased by ∼3.5-fold upon λ-phosphatase treatment relative to untreated conditions (Figure 6C). Figure S6D shows the phosphorylation levels of rACC1 alongside total rACC1 levels at various time points following λ-phosphatase treatment, demonstrating that λ-phosphatase efficiently dephosphorylates proteins without altering their total levels.

Inasmuch as each rACL, rACSS2, rACC1, and rFAS protein exhibited increased enzymatic activity in the dephosphorylated state, these observations alone could not distinguish whether the enhanced palmitate production resulted from cooperative complex formation or simply reflected additive increases in the intrinsic activities of individual enzymes upon dephosphorylation. To directly address this issue, we performed a FA-synthesis assay using (1) liver cytosol alone, (2) recombinant ACL–ACC1–FAS alone, and (3) a mixture of liver cytosol and recombinant ACL–ACC1–FAS, with protein amounts normalized across conditions. Each group was subjected to either dephosphorylated or phosphorylated conditions under identical treatment settings. As shown in Figure S6E, dephosphorylation was successfully achieved in all conditions. Under dephosphorylated conditions, combining liver cytosol with recombinant ACL–ACC1–FAS produced substantially greater palmitate than the expected additive activity calculated from cytosol alone and recombinant enzymes assayed in parallel (Figure 6D). This enhancement exceeded what would be predicted from simple additive effects of dephosphorylation on individual enzymes and was supported by a significant interaction between phosphorylation state and mixing condition (two-way ANOVA, F (3,16) = 133.4, P < 0.0001). Together, these data indicate a strong synergistic increase in lipogenesis arising from cooperative interactions between dephosphorylated cytosolic proteins and recombinant enzymes. In contrast, under phosphorylated conditions, the palmitate produced by the combined reaction was indistinguishable from the expected additive activity (Figure 6E), indicating that cooperative assembly and synergistic flux enhancement are selectively enabled by dephosphorylation.

To directly assess complex formation under these conditions, ACC1 was immunoprecipitated from the same reaction mixtures. Consistent with the results shown in Figure 5A, dephosphorylation markedly enhanced the interactions of ACC1 with FAS and ACL, and β-tubulin as determined by co-precipitation studies, suggesting that the lipogenic metabolon may be functionally linked to cytoskeletal components (Figure S6F). No nonspecific binding was observed with control rabbit IgG.

While it is not feasible to systematically test all cytosolic proteins that could potentially contribute to metabolon assembly, Figure 5 identified structural proteins as representative non-lipogenic factors that interact with ACC1 under dephosphorylated conditions. Consistent with this observation, β-tubulin showed detectable co-localization with ACC1 around lipid droplets in primary hepatocytes under lipogenic refeeding conditions (Pearson’s R ≈ 0.32), whereas this spatial association was markedly reduced upon fasting (Figure 6F). In parallel, pharmacological perturbation of microtubule dynamics provided support for the role of cytoskeleton in lipogenesis. Disrupting microtubule polymerization with nocodazole consistently decreased FA synthesis in primary hepatocytes, whereas stabilizing microtubules with taxol, an agent with the opposite effect of nocodazole, had minimal impact on lipogenesis (Figure 6G). These observations suggest that intact cytoskeletal architecture facilitates lipogenesis by coordinating the nutrient-responsive assembly and spatial organization of lipogenic enzyme complexes, rather than directly regulating microtubule dynamics.

Collectively, these findings suggest that lipogenic activity is enhanced through post-translational regulation that promotes the formation of a functional lipogenic metabolon under physiological conditions favoring high DNL. This process likely involves dynamic and cooperative interactions among DNL- and TG-related enzymes, aided by structural proteins and other accessory factors, allowing for more integrated and efficient lipogenesis.

### ACC1 anchors a multi-module lipogenic network integrating translation, trafficking, and lipid metabolism

To further map the ACC1-centered protein network, we immunoprecipitated ACC1 from wild-type and ACC-deficient liver lysates. In parallel, MIG12 and S14, two direct ACC1-binding partners were immunoprecipitated, as was ACSS2, a representative lipogenic enzyme, to compare their interaction profiles (Figure 7A). MIG12 and S14 pulldowns closely mirrored the ACC1 immunoprecipitation pattern: proteins enriched in wild-type lysates were markedly reduced in ACC-deficient lysates, consistent with their direct association with ACC1 and the dependence of their interactomes on ACC1 integrity. In contrast, ACSS2 immunoprecipitation showed a distinct profile. Although several lipogenic enzymes were reduced in ACC-deficient lysates, ACSS2 retained clear interactions with multiple lipogenic proteins. This pattern suggests that ACC1 enhances or stabilizes the overall lipogenic assembly rather than acting as its sole organizing component. Notably, β-actin binding to ACSS2 was unchanged regardless of ACC1 status, implying that actin contributes to complex organization through an ACC1-independent mechanism.

**Figure 7.**
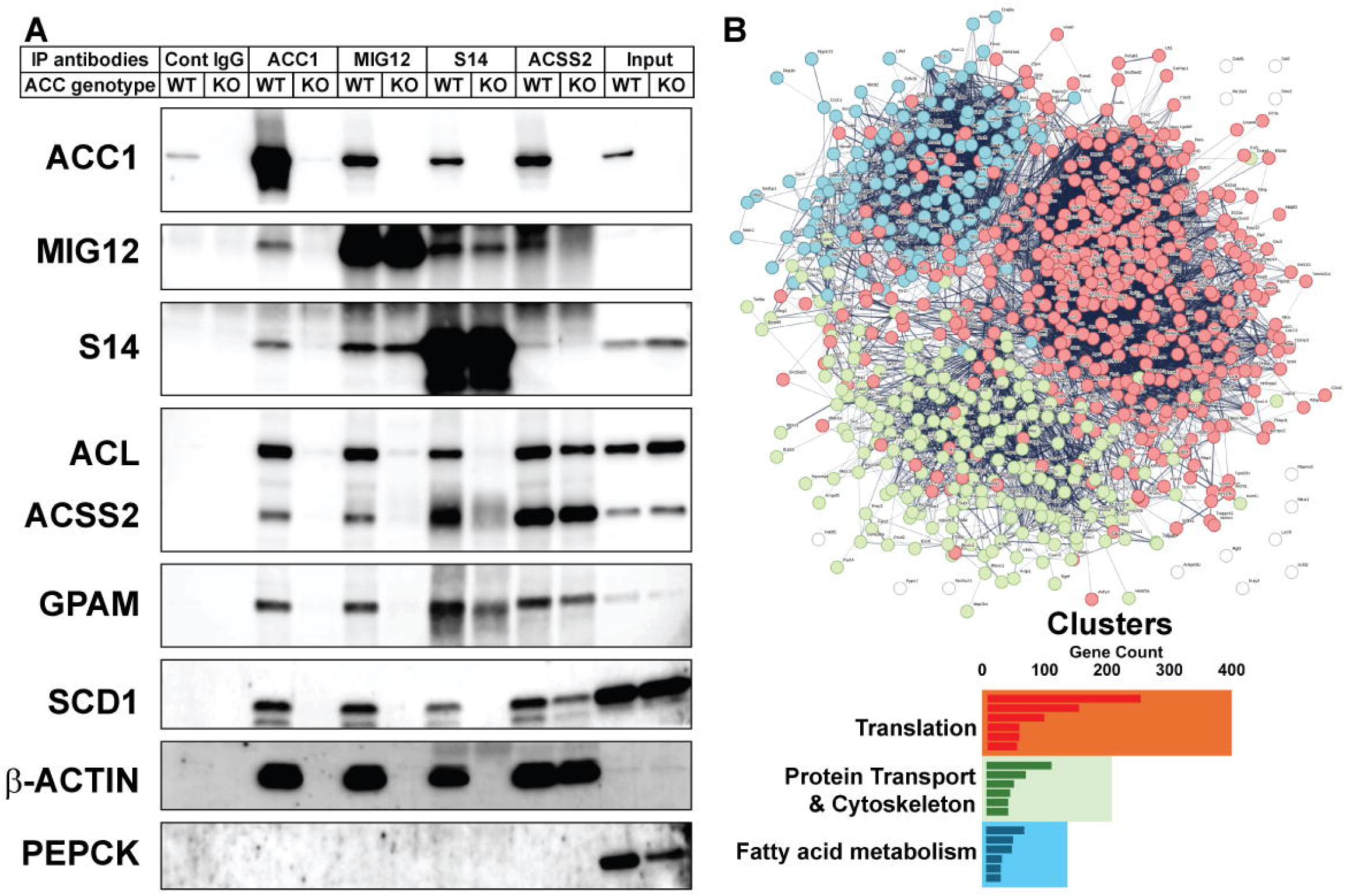
ACC1 anchors a multi-module lipogenic network integrating translation, trafficking, and lipid metabolism. (A) Liver lysates were prepared from *ob/ob* mouse livers and ACC1/ACC2–deleted *ob/ob* mouse livers as described in Methods. To activate endogenous phosphatases, lysates were incubated in 5 mM MgCl₂ for 30 minutes at 30°C. In parallel, anti-ACC1, anti-MIG12, anti-S14, or anti-ACSS2 antibodies were pre-bound to protein G magnetic beads by incubating for 30 minutes at room temperature. The activated lysates were then added directly to the antibody–bead complexes and incubated for 1 hour at room temperature to perform immunoprecipitation. After extensive washing, bound proteins were eluted by boiling and analyzed by SDS-PAGE and immunoblot analysis was carried out with the indicated antibodies. (B) Proteins co-immunoprecipitated with ACC1, MIG12, or S14 that were reduced by >20-fold in ACC-deficient lysates were classified as ACC1-dependent interactors and subjected to STRING network analysis. Three major functional clusters emerged—translation (red), protein trafficking/transport and cytoskeletal protein binding (green), and FA metabolism (blue). These clusters are displayed both as a network graph and summarized in the bar graph below using the corresponding color scheme. The inset mini-graphs within each colored bar summarize representative GO/KEGG terms for that cluster, with full enrichment analyses provided in Figure S7.

We next quantified proteins co-immunoprecipitated with ACC1 by mass spectrometry. Proteins that were reduced by >20-fold in ACC1, MIG12, or S14 immunoprecipitations from ACC-deficient lysates were classified as ACC1-dependent interactors. STRING network analysis of these proteins resolved three highly connected clusters (Figure 7B, top). The largest cluster consisted of ribosomal and translation-associated proteins, suggesting that a robust interface between ACC1 and the translational machinery exists. A second cluster was enriched for vesicle/trafficking components and cytoskeletal protein-binding factors, indicating that ACC1 is embedded within a broader transport and structural scaffold. A third cluster consisted of enzymes involved in FA synthesis and remodeling, consistent with ACC1’s canonical metabolic function (Figure 7B, bottom). DAVID functional annotations performed on the same dataset produced closely matching categories (Figure S7 and Supplemental Table 3).

Together, these findings indicate that activated ACC1 functions as a central organizational hub that coordinates core lipogenic enzymes—including FAS, ACL, ACSS2, SCD1, GPAM, and ACC2—into a coherent metabolic assembly. Direct binding by MIG12 and S14 likely stabilizes this structure, while cytoskeletal and trafficking components associate with the ACC1-centered complex to support spatial organization and substrate delivery. The presence of ribosomal proteins and transport-associated factors among ACC1-dependent interactors further suggests that lipogenesis is integrated with protein synthesis and intracellular trafficking through multiple points of connection.

## DISCUSSION

Transcriptional control of FA and TG synthesis is well characterized ^54^, but much less is known about how lipid synthesis is regulated post-transcriptionally. Here, we identified a previously unrecognized post-translational mechanism in which hepatic lipogenic enzymes assemble into a multi-protein metabolon under anabolic conditions. This metabolon provides a spatial basis for efficient glycerolipid synthesis and introduces a new dimension to the regulation of DNL.

Using immunofluorescence and EM, we showed that enzymes involved in FA and TG synthesis, along with accessory proteins like MIG12 and S14, co-localize around LDs and function as an integrated multi-enzyme assembly rather than as isolated enzymes under physiological activation states such as high carbohydrate feeding. These findings build on earlier LD proteomics reports that identified ACC1 ^55–59^ and FAS ^58^ on LDs, extending them by providing direct structural visualization of their co-localization in mammalian hepatocytes. Furthermore, the proximity between cytosolic enzymes such as ACC1 and FAS and the mitochondrial enzyme GPAM at contact sites suggests coordinated substrate transfer into the glycerolipid pathway. Subcellular fractionation of liver lysates from high-carbohydrate-refed rats further supports these observations.

Immunoprecipitation of ACC1 from liver lysates incubated under phosphatase-active conditions revealed robust interactions between ACC1 and multiple lipogenic proteins, whereas the inclusion of phosphatase inhibitors markedly reduced these associations (Figure 5A). This biochemical shift was paralleled by *in vitro* FA and TG synthesis assays, in which phosphatase-active conditions produced substantially higher lipid synthesis than phosphatase-inhibited ones. We initially attributed this enhanced interaction and activity to ACC1 dephosphorylation, a hypothesis supported by *in vivo* experiments using the AMPK agonist AICAR. In primary hepatocytes isolated from high-carbohydrate–refed rats, AICAR treatment caused the activated, polymerized form of ACC1 to dissociate into its inactive dimeric form, as shown by blue native PAGE. The extensive polymerization of ACC1 driven by MIG12 overexpression was also significantly reduced by AICAR-mediated ACC1 phosphorylation. Correspondingly, short-term AICAR treatment significantly suppressed hepatic lipogenesis in primary hepatocytes without affecting the steady-state levels of lipogenic enzymes and markedly reduced ACC1 localization around LDs. Notably, ACC1 was not the only enzyme influenced by dephosphorylation, as phosphatase treatment also increased the enzymatic activities of rACSS2 and rFAS, although to a lesser extent than ACC1. These findings suggest that anabolic phosphatase-active states coordinate multiple components of the lipogenic pathway, collectively enhancing metabolon assembly and catalytic output.

Another notable finding from the immunoprecipitation experiments was the strong interaction of ACC1 with ACC2 and MIG12. Whereas most ACC1-associated proteins showed enhanced binding under phosphatase-active conditions, ACC2 and MIG12 maintained stable interactions with ACC1 irrespective of its phosphorylation state. This feature suggests that these two proteins form a relatively stable core structural interaction with ACC1.

Additional insight into ACC2 was obtained from transfection studies in CHO-K1 cells. When ACC1 and ACC2 were co-expressed, the fraction of ACC1 that interacted with ACC2 and formed heterodimers or higher-order heteromultimers and localized to mitochondria increased (Figure S5D). ACC1 also retained a substantial cytosolic pool that did not overlap with ACC2, likely reflecting cytosolic homomeric ACC1 assemblies. These observations indicate that ACC1 is partitioned between heteromeric ACC1–ACC2 complexes at mitochondria and homomeric ACC1 assemblies in the cytosol and suggest that the subcellular localization and functional partitioning of malonyl-CoA–producing acetyl-CoA carboxylase activity may be influenced by the relative abundance of ACC2. These findings are consistent with the tissue-specific expression patterns of ACC isoforms. In liver and adipose tissue, where malonyl-CoA is used primarily for lipogenesis, ACC1 is the predominant isoform; whereas in muscle and heart, where malonyl-CoA mainly inhibits mitochondrial fatty acid oxidation, ACC2 is the major form. Thus, cells may regulate the relative abundance (stoichiometry) of ACC1 and ACC2 to redistribute acetyl-CoA carboxylase activity between the cytosol and mitochondria according to metabolic demands. In this context, ACC1–ACC2 interactions may serve as a regulatory node linking FA synthesis and oxidation. Alterations in the stoichiometry of the two isoforms could therefore shift the balance between lipogenesis and control of mitochondrial FA oxidation by modulating the subcellular distribution of carboxylase activity.

Finally, although ACC2 has long been viewed primarily as a regulator of mitochondrial FA oxidation, ACC2 deletion reduced ACC1 interactions with multiple lipogenic enzymes and structural proteins (Figure 5B). This finding supports the view that ACC2 functions not only as a metabolic enzyme that inhibits FA oxidation but also as a structural component essential for the organization and maintenance of the lipogenic metabolon. Complementing these structural insights, our biochemical assays demonstrated that assembling lipogenic enzymes within a shared reaction environment generates catalytic outputs that exceed the sum of individual activities. Under DNL-activated conditions, combining recombinant ACL–ACC1–FAS with liver cytosol nearly doubled the FA synthesis compared with the expected additive activity, indicating synergistic enhancement of pathway flux. This synergy suggests that enzyme co-assembly, rather than simply increased intrinsic activity, facilitates more efficient substrate channeling during active lipogenesis. These functional observations reinforce the multi-module organization described in Figure 7, in which ACC1 immunocomplexes interact not only with lipogenic enzymes but also with ER-proximal translation components, vesicle-movement proteins, and actin-linked scaffolds, supporting the concept that lipogenic enzymes achieve maximal efficiency when spatially integrated with cytoskeletal scaffolds and intracellular transport machinery, and when functionally linked to translation-associated components.

Although metabolon-based regulation is less explored in lipogenesis, analogous mechanisms have been established in the TCA cycle and glycolysis ^29–31,60–63^. The substrate-responsive assembly of PFK filaments in the presence of fructose-6-phosphate ^64^ provides a relevant precedent and parallels ACC1 filament formation under anabolic conditions, suggesting that these filaments may serve as structural scaffolds for metabolon assembly. Consistent with this concept, a FA–synthesis metabolon has been described in the thermophile *Chaetomium thermophilum* ^65^ and ACC–FAS co-localization has been visualized in *Saccharomyces cerevisiae* ^66^, where these enzymes are reorganized or sequestered in response to glucose starvation. These observations across lower eukaryotes support the idea that nutrient state governs the assembly and disassembly of spatially organized lipogenic enzymes.

Recent work by Wijesinghe *et al.* ^67^ extends this concept to mammalian cells. In their study, ACC1 formed ring-like puncta and localized to the surface of lipid droplets (LDs), but this localization was highly conditional: anabolic cues such as AMPK inhibition preserved ACC1–LD co-localization, whereas large LDs generated by acute oleic-acid loading—which diminishes anabolic drive—displayed reduced ACC1 staining. This pattern mirrors our observation that the AMPK agonist AICAR suppresses ACC1 accumulation around LDs (Figure 4E), indicating that ACC1 recruitment to LDs requires appropriate metabolic activation rather than merely reflecting LD abundance. Together, these findings align with our model in which ACC1 polymerization and its interactions with accessory factors generate a nutrient-responsive lipogenic complex in hepatocytes. Collectively, the emerging evidence across species supports a broader framework in which ACC1 integrates nutrient status with spatial enzyme organization to optimize lipogenesis during anabolic transitions.

In conclusion, our study demonstrates that under anabolic conditions, hepatic lipogenic enzymes assemble into a spatially organized, phosphorylation-responsive metabolon around LDs. This multi-protein metabolon enhances catalytic efficiency facilitating improved transfer of intermediates between sequential enzymes, thereby supporting efficient synthesis of FA and TGs. These findings provide a mechanistic foundation for understanding hepatic lipid synthesis and highlight therapeutic opportunities in metabolic disease by targeting the structure and regulation of the lipogenic metabolon.

### Limitations of the study

In this study, we present evidence for a novel post-translational regulatory mechanism that enhances FA and TG synthesis in the liver via the assembly of a multi-protein metabolon. Ideally, since the structure of ACC1 under different activation states has been reported ^68,69^, we would have included structural analyses of this ACC1 mediated lipogenic metabolon. However, detecting such metabolons presents greater challenges compared to stable protein assemblies. The reason is that a metabolon is characterized as a transient structural-functional metabolon formed by sequential enzymes within a metabolic pathway, exhibiting non-covalent interactions and associating with various cellular structural elements, such as integral membrane proteins and cytoskeletal components ^30,70^. Consequently, the interactions among these proteins are more transient and dynamic, enabling them to adapt to fluctuations in metabolic environments.

Additionally, the mRNA levels of the lipogenic proteins are all significantly downregulated during fasting as a result of reduced SREBP-1c expression, which restricted our ability to compare the co-localization of lipogenic proteins between fasting and refeeding conditions. Therefore, we employed AICAR treatment and elevated phosphorylation levels to simulate catabolic states *in vivo* and *in vitro*, respectively.

### Declaration of generative AI and AI-assisted technologies in the writing process

During the preparation of this manuscript, the authors used ChatGPT (OpenAI) solely for language editing, including grammar correction and improvement of clarity. No content, data interpretation, or scientific conclusions were generated by the tool. After using this assistance, the authors thoroughly reviewed, revised, and approved all text, and they take full responsibility for the integrity and accuracy of the final manuscript.

## RESOURCE AVAILABILITY

### Lead contact

Further information and requests for resources should be directed to the lead contact, Jay D. Horton.

### Materials availability

Custom antibodies generated in this study are available upon reasonable request and completion of an MTA. Cloning constructs are available from the Lead Contact. All other materials used in this study are specified in the key resources table.

### Data and code availability

All data supporting the findings are available within the paper and Supplemental Information. No custom code was used.

## ACKNOWLEDGEMENTS

We thank Norma Anderson, Judy Sanchez, Tuyet Dang, and Tessa Edwards for excellent technical assistance; Lisa Beaty for suspension cell culture and set up; Linda Donnelly for assistance with antibody generation; Jinhai Yu for providing TEV protease; and Chelsea Burroughs and Nancy Heard for help with graphics. The work was supported by grants NIH P01HL160487 and NIH P30DK127984.

## AUTHOR CONTRIBUTIONS

X.Z., W.B.M., T.T., X.F., and C.-W.K. conducted the experiments. J.D.H., C.-W.K., and X.Z. designed the experiments. X.Z., C.-W.K., and J.D.H. wrote the manuscript. All authors reviewed and approved the article.

## Key resource table

| REAGENT or RESOURCE | SOURCE | IDENTIFIER |
| --- | --- | --- |
| **Antibodies** |  |  |
| Rabbit anti-ACL | Abcam | Cat# ab40793 |
| Rabbit anti-ACSS2 | Abcam | Cat# ab66038 |
| Rabbit anti-mouse ACC1 | House antibodies<sup>71</sup> | 508D or 510D |
| Mouse monoclonal anti-mouse ACC1 | House antibodies | 2B5 |
| Rabbit anti-mouse ACC2 | House antibodies | 456D |
| Rabbit anti-human FAS | House antibodies | 310D |
| Rabbit anti-mouse MIG12 | House antibodies<sup>72</sup> | 143D |
| Mouse monoclonal anti-mouse MIG12 | House antibodies | 6C4 |
| Rabbit anti-mouse S14 | House antibodies | 39C |
| Rabbit anti-PEPCK | Abcam | Cat# ab70358 |
| Mouse anti-β-Tubulin | ThermoFisher | Cat# MA5-16308 |
| Rabbit anti-β-Actin | Cell Signaling | Cat# 4970S |
| Rabbit Anti-mouse GPAM | House antibodies | 280D |
| Mouse monoclonal anti-rat GPAM | House antibodies | 16F8 |
| Anti-ADRP | Sigma | Cat# SAB4200452 |
| Anti-SCD1 | Cell Signaling | Cat# 2794S |
| Anti-PROHIBITIN | Santa Cruz | Cat# SC-56467 |
| Anti-CYTOCHROME C | Millipore | Cat# MAC1800 |
| Mouse anti-CPT1 | Abcam | Cat# ab128568 |
| Anti-Ser79-pACC1 | Cell Signaling | Cat# 3661S |
| Anti-PDI | Cell Signaling | Cat# 2446S |
| Goat Anti-Rabbit IgG, F(ab')<sub>2</sub> | Jackson Immuno | Cat# 111-035-047 |
| Goat Anti-Mouse IgG, F(ab')<sub>2</sub> | Jackson Immuno | Cat# 115-035-072 |
| Li-COR IRDye 800 goat anti-Rabbit IgG | Fisher Scientific | Cat# NC9401842 |
| Li-COR IRDye 680 goat anti-Mouse IgG | Fisher Scientific | Cat# NC0252290 |
| Li-COR IRDye 680RD Streptavidin | Fisher Scientific | Cat# NC0337633 |
| **Bacterial and virus strains** |  |  |
| Stellar™ Competent Cells | Takara | Cat# 636764 |
| TOP10 Competent Cells | ThermoFisher | Cat# C404003 |
| **Chemicals, peptides, and recombinant proteins** |  |  |
| LB broth | Research Products International | Cat# L24061 |
| Freestyle 293 expression medium | Gibco | Cat# 12338018 |
| Opti-MEM™ I Reduced Serum Medium | Gibco | Cat# 31985070 |
| Fetal Calf Serum (FCS) | Sigma-Aldrich | Cat# F2442 |
| Penicillin-Streptomycin solution (100x) | Corning | Cat# 30-002-CI |
| PEI 25K | Polysciences | Cat# 23966-1 |
| Protease Inhibitor Cocktail Set III, EDTA-Free - Calbiochem | Millipore-Sigma | Cat# 539134-1ML |
| Protease Inhibitor Cocktail, for use in purification of Histidine-tagged proteins, DMSO solution | Sigma-Aldrich | Cat# P8849-5ML |
| Purified rTEV protease | Gift from Dr. Jinhai Yu, UT Southwestern Medical Center | N/A |
| Lambda Protein Phosphatase | New England Biolabs | Cat# P0753L |
| DL-Dithiothreitol | Sigma-Aldrich | Cat# 43819-5G |
| Adenosine 5'-triphosphate magnesium salt | Sigma-Aldrich | Cat# A9187-1G |
| Adenosine 5'-triphosphate disodium salt hydrate | Sigma-Aldrich | Cat# A2383-5G |
| Coenzyme A sodium salt hydrate | Sigma-Aldrich | Cat# C3144-25MG |
| Acetyl coenzyme A sodium salt | Sigma-Aldrich | Cat# A2056-25MG |
| Malonyl coenzyme A lithium salt | Sigma-Aldrich | Cat# M4263-10MG |
| NADPH, Tetrasodium Salt | Sigma-Aldrich | Cat# 481973-100MG |
| Sodium bicarbonate [14C] | American Radiolabeled Chemicals | Cat# ARC 0138C-50 µCi |
| Malonyl [2-14C] Coenzyme A | American Radiolabeled Chemicals | Cat# ARC 0528-25 µCi |
| Citric Acid, [1,5-14C] | Revvity | Cat# NEC160050UC |
| Acetic Acid, Sodium Salt, [1,2-14C] | Revvity | Cat# NEC553050UC |
| Lauryl Maltose Neopentyl Glycol | Anatrace | Cat# NG310 5 GM |
| 3a70B Complete Counting Cocktail | Research Products International | Cat# 111154 |
| In-Fusion® Snap Assembly Value Bundle | Takara | Cat# 638946 |
| Platinum™ SuperFi II PCR Master Mix | ThermoFisher | Cat# 12368010 |
| DreamTaq Green PCR Master Mix (2X) | ThermoFisher | Cat# K1081 |
| Oleoyl coenzyme A dipotassium salt | ThermoFisher | Cat# CN10090405 |
| sn-Glycerol 3-phosphate lithium salt | Sigma-Aldrich | Cat# 94124-10MG |
| Glycerol, L-[14C(U)] 3-phosphate disodium salt | American Radiolabeled Chemicals | Cat# ARC 0316-50 µCi |
| Sodium butyrate | Sigma-Aldrich | Cat# 303410-500G |
| HCS LipidTOX™ Deep Red Neutral Lipid Stain, for cellular imaging | Fisher Scientific | Cat# H34477 |
| Clean-Blot™ IP Detection Reagent (HRP) | Thermo Scientific | Cat# 21230 |
| AICAR | Abcam | Cat# ab120358 |
| **Critical commercial assays** |  |  |
| Pierce™ Coomassie (Bradford) Protein Assay Kit | ThermoFisher | Cat# 23200 |
| Pierce BCA Protein Assay Kit | ThermoFisher | Cat# 23225 |
| Silver staining Kit | ThermoFisher | Cat# 24612 |
| **Experimental models: Cell lines** |  |  |
| HEK-293S GnT1<sup>-</sup> | ATCC | Cat# CRL-3022;<br>RRID: CVCL_A785 |
| CHO-K1 | ATCC | Cat# CCL-61;<br>RRID: CVCL_0214 |
| **Experimental models: Organisms/strains** |  |  |
| C57BL/6J mouse | Jackson Laboratory | Stock# 000664 |
| ob/ob mouse | Jackson Laboratory | Stock# 000632 |
| Sprague Dawley | Charles River | SAS SD code 400 |
| **Phosphatase inhibitor cocktail** |  |  |
| Sodium Fluoride | Sigma-Aldrich | Cat# S7920 |
| Imidazole | Sigma-Aldrich | Cat# I5513 |
| Sodium molybdate | Sigma-Aldrich | Cat# M1003 |
| Sodium tartrate dihydrate | Sigma-Aldrich | Cat# S4797 |
| Sodium pyrophosphate | Sigma-Aldrich | Cat# P8135 |
| β-glycerophosphate | Sigma-Aldrich | Cat# G9422 |
| **Recombinant DNA** |  |  |
| pEZT-BM | Addgene | Plasmid #74099 |
| ATP citrate lyase (ACLY) (NM_001096) Human Tagged ORF Clone | Origene | Cat# RC200508 |
| ACSS2 (NM_018677.4) cDNA ORF clone, Homo sapiens (human) | Genscript | Cat# OHu18043 |
| ACL-Tev-Flag-His-XT3 | IDT DNA | N/A |
| ACC1-Tev-Flag-His-XT3 | IDT DNA | N/A |
| FAS-Tev-Flag-His-XT3 | IDT DNA | N/A |
| MIG12-Tev-Flag-His_Human | IDT DNA | N/A |
| S14-Tev-Flag-His_Human | IDT DNA | N/A |
| PEPCK_Human-Tev-His | IDT DNA | N/A |
| pEZT-BM-ACL-3xFlag-10xHis | This paper | N/A |
| pEZT-BM-ACSS2-3xFlag-10xHis | This paper | N/A |
| pEZT-BM-ACC1-3xFlag-10xHis | This paper | N/A |
| pEZT-BM-FAS-3xFlag-10xHis | This paper | N/A |
| pEZT-BM-MIG12-3xFlag-10xHis | This paper | N/A |
| pEZT-BM-S14-3xFlag-10xHis | This paper | N/A |
| pEZT-BM-PEPCK-3xFlag-10xHis | This paper | N/A |
| ACC1-pcDNA3.1 | This paper | N/A |
| ACC2-pcDNA3.1 | This paper | N/A |
| **Software and algorithms** |  |  |
| Image Studio™ Version 5.0 | Li-COR | N/A |
| Prism 10 | GraphPad | https://www.graphpad.com/ |
| Adobe Illustrator 2023 | Adobe | N/A |
| **Other** |  |  |
| PD-10 desalting columns | Cytiva | Cat# 17085101 |
| Superose 6 10/300 GL column | Cytiva | Cat# GE17-5172-01 |
| Amicon Ultra centrifugal filter | Millipore | Cat# UFC903008 |
| Ni-NTA Resin | Thermo Scientific | Cat# 88221 |
| Slide-A-Lyzer dialysis cassette | Thermo Scientific | Cat# 66203 |
| Dynabeads Protein A beads | Invitrogen | Cat# 10002D |
| Scintillation vials | Research Products International | Cat# 121040 |
| Disposable Round-Bottom Rimless Glass Tubes | Corning | Cat# 14-962-15E |
| 4–15% Criterion™ TGX™ Precast Midi Protein Gel, 26 well, 15 µl | BIO-RAD | Cat# 5671085 |
| 4–15% Mini-PROTEAN® TGX™ Precast Protein Gels, 15-well, 15 µl | BIO-RAD | Cat# 4561086 |
| Dynabeads™ Protein G for Immunoprecipitation | Fisher Scientific | Cat# 10-004-D |
| Slide Warmer (110 V, for 23 slides) | Fisher Scientific | Cat# 50-930-986 |

## Methods

### Experimental model and study participant details

#### Animals and primary hepatocytes

Primary hepatocytes were isolated from 4-week-old male Sprague–Dawley rats fasted for 12 hours and refed for 12 hours with a fat-free diet, using collagenase perfusion as described by Shimomura et al. ^73^ All rat studies were performed under an approved protocol from the UT Southwestern Institutional Animal Care and Research Advisory Committee (IACUC #2015-100838). Liver tissues were collected from wild-type male C57BL/6 mice, liver-specific ACC1 and/or ACC2 knockout male C57BL/6 mice, or ob/ob male C57BL/6 mice. All mice were maintained under standard housing conditions with a 12-hour light/dark cycle (lights on at 6 AM and off at 6 PM). Mouse studies were conducted under UT Southwestern IACUC approval (2015-101135-G). Nocodazole and taxol were used at 20 μM to achieve robust disruption or stabilization of microtubules, respectively, in primary hepatocytes.

### Method Details

#### Antibody generation

For the generation of the mouse monoclonal antibody that recognizes mouse ACC1, a cDNA encoding the N-terminal 95 amino acids of mouse ACC1 was cloned into a pGEX-4T-1 vector and the recombinant protein was expressed in *Escherichia coli (E. coli)* and purified using glutathione agarose. Monoclonal antibodies were generated by fusion of Sp2/mIL-6 (CRL-2016; ATCC) mouse myeloma cells with splenic B lymphocytes derived from a female NZBWF1/J mouse injected with purified mouse ACC1 protein ^74,75^.

For the generation of a rabbit polyclonal antibody against mouse ACC2, a cDNA encoding the N-terminal 200 amino acids of mouse ACC2 was cloned into a pRSF-1b expression vector and expressed in *E. coli*. The purified His-tagged protein was isolated using nickel affinity resin and injected into rabbits as described ^76^.

For the generation of rabbit polyclonal antibody against human FAS, a cDNA encoding the C-terminal 305 amino acids from human FAS was cloned into pGEX-4T-1 expression vector and expressed in *E. coli.* The purified GST-fusion protein was injected into rabbits as described ^19^.

For the generation of a rabbit polyclonal antibody against mouse S14, recombinant (His)6-mouse S14 was expressed in E. coli, purified by Ni-NTA affinity chromatography (Qiagen) according to the manufacturer’s protocol, and injected into rabbits as described ^19^.

For the generation of rabbit polyclonal antibody against mouse GPAM, a cDNA encoding the N-terminal 150 amino acids from mouse GPAM was cloned into pRSF-1b expression and expressed in *E. coli*. The purified His-tagged protein was injected into rabbits as described ^19^.

For the generation of mouse monoclonal antibody against rat GPAM, a cDNA encoding amino acids 627 to 717 of rat GPAM was cloned into pRSF-1b expression vector and expressed in *E. coli*. The purified His-tagged protein was injected into mice, and monoclonal antibodies were produced as described above. For the generation of mouse monoclonal antibody against mouse MIG12, full length mouse MIG12 cDNA was cloned into pRSF-1b and expressed in *E. coli* and the purified protein (Ni-NTA agarose) was injected into mice. Monoclonal antibodies were generated using the same hybridoma fusion strategy described for ACC1.

#### Immunofluorescence

Primary hepatocytes from refed rat livers were fixed in 4% paraformaldehyde (PFA) in PHEM buffer (pH 7.4) for 30 minutes. After rinsing the cells with 0.1 M phosphate buffer, they were permeabilized with pre-chilled (−20°C) methanol for 15 min at −20°C. Blocking was performed using 1% BSA and 0.2% fish skin gelatin in 0.1 M phosphate buffer (pH 7.4) for 30 minutes at room temperature. Primary antibodies were incubated overnight at 4°C in the blocking solution, and Alexa Fluor 488- or 568-labeled secondary antibodies were used to visualize the localization of each target protein. Imaging was conducted using a Zeiss LSM880 Airyscan microscope.

#### Colocalization analysis

Colocalization was quantified using the Coloc2 plugin in Fiji/ImageJ. To minimize background noise and evaluate biologically relevant signal overlap, analysis was performed on lipid droplet–enriched regions, which represent functional sites of lipogenic enzyme accumulation. CZI files were imported directly into Fiji, and channels were split and processed separately to avoid spectral bleed-through. Images used for Coloc2 analysis were converted to 8-bit format prior to computation. Pearson’s correlation coefficients were calculated using Costes automatic thresholding, and above-threshold Pearson’s R values were reported in the main text. Thresholded Manders’ coefficients (tM1 and tM2), Costes significance testing, and 2D intensity histograms were generated by Coloc2 and are presented in Figure S1. All images were acquired below saturation, identical ROIs were applied to paired channels for each analysis.

#### Pre-embedding immuno-electron microscopy

Primary hepatocytes from refed rat livers were fixed in 4% PFA/0.05% glutaraldehyde (GA) in PHEM buffer (pH 7.4) for 30 minutes. After rinsing the cells with 0.1 M phosphate buffer, residual aldehydes were quenched with 50 mM glycine in 0.1 M phosphate buffer for 15 minutes, followed by permeabilization with 0.2 mg/mL saponin for 30 minutes at room temperature. Blocking was performed using 5 μg/mL saponin, 1% BSA, 0.2% fish skin gelatin, and 5% goat serum in 0.1 M phosphate buffer (pH 7.4) for 15 minutes at room temperature. The primary antibody was incubated overnight at 4°C in the blocking solution. After incubation, cells were labeled with an anti-rabbit or anti-mouse IgG secondary antibody conjugated to 1.4 nm gold. Following post-fixation, the samples were subjected to gold enhancement, embedded in Embed-812 resin, and cut into 60 nm sections. Electron micrographs were taken using the TEM JEOL 1400 Plus microscope. For ACC1 and FAS double immunogold labeling, ACC1 and FAS primary antibodies were bound to 5 nm and 10 nm colloidal gold-conjugated secondary antibodies, respectively. After post-fixation, the cells were embedded in Embed-812 resin, sectioned into 60 nm slices, and imaged using the JEOL 1400 Plus Transmission Electron Microscope. The pre-embedding method followed the procedure described by Yamamoto *et al*. ^77^

#### Liver subcellular fractionation

Fresh rat or mouse liver was homogenized in 50 mM Tris (pH 7.4) and 250 mM sucrose in the presence of protease inhibitors. For lipid droplet isolation, half of the homogenate was transferred to a new tube, and buffer A (20 mM HEPES, pH 7.4, 100 mM KCl, 2 mM MgCl₂) was gently added to the top. The sample was centrifuged at 2,000 × g for 30 minutes, and the floating lipid layer was transferred to a new tube. This lipid layer was mixed with an equal volume of buffer A-500S (20 mM HEPES, pH 7.4, 100 mM KCl, 2 mM MgCl₂, and 500 mM sucrose) and gently overlaid with buffer A. This step was repeated twice to isolate a pure lipid layer. A glass pipette was used during lipid layer purification to minimize loss of lipid. Proteins from the lipid layer were extracted using an organic solvent as described ^78^ and resuspended in SDS-lysis buffer (50 mM Tris, pH 6.8, 50 mM NaCl, 1% SDS). For mitochondria and microsome fractionations, the remaining half of the liver homogenate was centrifuged at 1,000 × g for 10 minutes to remove the nuclear fraction. The supernatant was centrifuged at 10,000 × g for 20 minutes to pellet the mitochondrial fraction. The mitochondrial pellet was resuspended and washed twice more with homogenization buffer by repeated centrifugation at 10,000 × g for 20 minutes. The final mitochondrial pellet was resuspended in SDS-lysis buffer. The supernatant from the 10,000 × g centrifugation was further centrifuged at 100,000 × g for 30 minutes to sediment the microsomal fraction, which contains the endoplasmic reticulum. The microsomal pellet was rinsed with homogenization buffer and resuspended in SDS-lysis buffer.

#### Recombinant protein expression and purification

##### Expression

The cDNAs encoding human ACSS2, ACL, ACC1, FAS, MIG12, S14, and PEPCK were cloned into the pEZT-BM vector containing a TEV protease cleavage site, an SSG linker, followed by a C-terminal 3 × Flag tag and 10 × His tag. All proteins were expressed in mammalian HEK 293S GnTI⁻ cells (ATCC). A total of 500 mL of HEK 293S GnTI⁻ cells were transfected at a density of 2–3 × 10⁶ cells/mL with 0.75 mg of plasmid DNA using PEI 25K (Polysciences). 16 hours post-transfection, 250 mL of Freestyle 293 expression medium (Gibco) supplemented with 2% fetal calf serum (FCS) and 1% penicillin-streptomycin solution (Corning) was added to the cultures. Sodium butyrate was added to a final concentration of 8 mM to enhance protein expression. Cells were harvested 72 hours post-transfection for subsequent protein purification.

##### Purification

The cell pellet from 400 mL culture was resuspended and homogenized in 100 mL buffer B (50 mM phosphate buffer, 500 mM NaCl, 1 mM DTT, 5 mM imidazole, 225 mM mannitol, 75 mM sucrose, protease inhibitor cocktail (Millipore-Sigma), protease inhibitor cocktail for use in purification of Histidine-tagged proteins (Sigma-Aldrich), pH 8.0). The homogenate was centrifuged at 135,000 × g for 45 minutes at 4 °C. The supernatant was collected and filtered through a 5 μm filter. The filtered supernatant was loaded on a 2 mL Ni-NTA resin (Thermo Scientific) that pre-equilibrated with buffer B and flowed through by gravity (∼1 ml/min) at 4°C. The Ni-NTA was then washed sequentially with 10 CV buffer B and 25 CV buffer C (50 mM phosphate buffer, 500 mM NaCl, 1 mM DTT, 50 mM imidazole, 10% glycerol, pH 8.0) by gravity. Proteins were eluted in 5 CV buffer D (50 mM phosphate buffer, 500 mM NaCl, 1 mM DTT, 500 mM Imidazole, 10% glycerol, pH 8.0). The eluate was desalted using a PD-10 desalting column (Cytiva) pre-equilibrated with buffer E (10 mM HEPES, 50 mM NaCl, 100 mM KCl, 1 mM DTT, 5% glycerol, pH 7.4). Proteins were eluted in buffer E and incubated with TEV protease overnight at 4 °C.

##### Removal of TEV and tags

rACL, rACC1, and rFAS: the TEV-cleaved proteins were concentrated to 0.5 mL using a 30,000 MWCO Amicon Ultra centrifugal filter (Millipore) and then subjected to gel-filtration chromatography on a Superose 6 10/300 GL column (Cytiva) pre-equilibrated with buffer E. Purified proteins eluted as a single sharp peak in buffer E. Protein homogeneity was confirmed by Coomassie blue-stained SDS-PAGE.

rACSS2, rMIG12, rS14, and rPEPCK: the TEV-cleaved proteins were loaded on 2 mL Ni-NTA resin and flowed through by gravity (∼1 ml/min) pre-equilibrated with buffer E. The Ni-NTA resin was then washed sequentially with 5 CV buffer B, 5 CV buffer C and 5 CV buffer D by gravity flow. The flow-through and eluted fractions without His tag signal were combined and concentrated using an Amicon Ultra centrifugal filter (Millipore) (rACSS2 and rPEPCK: 30,000 MWCO, rMIG12: 10,000 MWCO, rS14: 3,000 MWCO) and then dialyzed using a 2,000 MWCO Slide-A-Lyzer dialysis cassette (Thermo Scientific) in 2 L buffer E overnight at 4°C.

##### Co-immunoprecipitation of recombinant proteins

Recombinant ACC1, FAS, ACL, ACSS2, MIG12, S14, and PEPCK were mixed at 8: 64: 20: 40: 32: 96: 1 molar ratios, respectively. Groups lacking recombinant ACC1 served as controls. The protein mixtures were incubated with 40 mM HEPES (pH 7.1), 1 mM MnCl₂, and λ-phosphatase (New England Biolabs) for 30 minutes at 30°C. For phosphorylated groups, the mixtures were incubated with a phosphatase inhibitor cocktail instead of lambda phosphatase, under the same conditions. The reaction systems were then supplemented with 0.5 mM Na-citrate buffer (pH 7.1) and incubated for 20 minutes at 30°C.

After pre-incubation, 5 μg of ACC1 polyclonal antibody was added to each reaction and incubated for an additional 10 minutes at 30°C. Buffer F (50 mM NaCl, 100 mM KCl, 20 mM HEPES, 4 mM MgCl₂, 1% LMNG, pH 7.1) was added, and the mixtures were incubated with rotation for 1 hour at room temperature. Subsequently, 25 μL Dynabeads Protein A (Invitrogen) were added and incubated with rotation for 20 minutes at room temperature. The beads were washed four times with buffer F. Bound proteins were eluted by incubation with 1× SDS sample buffer for 10 minutes at 95°C. Eluted proteins were analyzed by immunoblot analysis using the indicated antibodies.

##### Co-immunoprecipitation of liver lysates

Liver slices from fresh mouse (or rat) livers were homogenized in 10 mM Tris (pH 7.1), 250 mM sucrose, and 1% LMNG (Lauryl maltose neopentyl glycol) in the presence of protease inhibitors. The homogenate was centrifuged at 25,600 × g for 15 minutes at 20°C. The supernatant was collected and filtered through a 0.45 μm filter. The clarified supernatant was divided into two tubes; 5 mM MgCl₂ was added to both, whereas phosphatase inhibitor cocktail was added to only one. After incubation at 30°C for 30 minutes, NaCl and KCl were added to final concentrations of 50 mM and 100 mM, respectively. Mouse monoclonal ACC1 antibody or mouse IgG control antibody was added to each tube, and the samples were incubated for 1 hour at room temperature. Dynabeads Protein G (25 μL) were then added and incubated for a further 25 minutes at room temperature. The beads were washed 5 times with 10 mM Tris (pH 7.1), 100 mM KCl, 50 mM NaCl, 5 mM MgCl₂, and 1% LMNG. The beads were boiled in SDS-loading buffer for 5 minutes to elute proteins co-immunoprecipitated with ACC1.

##### Mass Spectrometry

Immunoprecipitates were resolved by SDS–PAGE, Coomassie-stained, and the excised gel regions were submitted to the UT Southwestern Proteomics Core Facility for LC–MS/MS analysis. Protein identification and label-free quantification were performed using the core’s standard Complex Mixture ID workflow, and processed datasets were used for downstream analyses.

##### Radiometric FA synthesis in hepatocytes

Rat primary hepatocytes were cultured in 6-well dishes coated with collagen. Where indicated, after 8 hours of culture, cells were treated with DMSO, nocodazole, or taxol (20 μM). On the day of study, the culture medium was replaced with medium containing 1 mM acetate or citrate, supplemented with 1 μCi [¹⁴C]-labeled acetate or citrate, respectively, and cells were harvested after the indicated incubation times by adding 1 mL of 0.1 N NaOH. A recovery carrier (^3^H oleic acid), 1.5 mL of ethanol, and 0.75 mL of 75% KOH were added to the harvested cells, and the mixture was autoclaved for 45 minutes on slow exhaust for saponification. After saponification, 1.5 mL of ethanol and 1.5 mL of HCl were added sequentially with vigorous vortexing. After cooling, 1.5 mL of ethanol was added, and fatty acids were extracted three times with 3 mL of hexane. The solvent was evaporated under nitrogen, and the residue was re-dissolved in 30 μL of chloroform. Samples were spotted onto plastic-backed silica gel TLC plates and resolved in a solvent system consisting of heptanes:diethyl ether:acetic acid (90:30:1, v/v/v). After visualization with iodine vapor, each FA spot was excised and radioactivity was quantified by scintillation counting. FA synthesis was calculated based on the recovered radioactivity.

#### Recombinant protein enzyme assays

##### Dephosphorylation treatment

Different recombinant proteins were pre-incubated with 20 mM HEPES (pH 7.1), 1 mM MnCl_2_, and lambda phosphatase for 30 minutes at 30°C in 96 well plates (Corning). The same groups without lambda phosphatase were used as phosphorylated groups.

##### ACSS2 and ACL (LC-MS/MS)

For the ACL activity assay, protein samples from dephosphorylation treatment were incubated at 37°C in a reaction mixture containing 20 mM HEPES (pH 7.1), 1 mM DTT, 1 mg/mL BSA, 4 mM sodium citrate (pH 7.1), 5 mM Mg-ATP (pH 7.1), and 0.1 mM CoA. For the ACSS2 activity assay, protein samples were incubated at 37°C with 20 mM HEPES (pH 7.1), 1 mM DTT, 1 mg/mL BSA, 4 mM sodium acetate (pH 7.1), 5 mM Mg-ATP (pH 7.1), and 0.1 mM CoA. At designated time points (0, 4, 8, 16, and 32 minutes), reactions were terminated by the addition of 8 N HCl, followed by thorough mixing. Each sample was then immediately spiked with a [1-¹³C]acetyl-CoA internal standard prior to further analysis.

Samples were centrifuged at 4°C for 10 minutes and 150 μL supernatant was loaded on an Oasis HLB 1cc-30 mg solid-phase extraction column. The column was washed with water and methanol. The eluent was lyophilized and dissolved in 50 μL of mobile phase of which 1 μL was subjected to LC/MS/MS analyses. Analysis was performed on an API 3200 triple quadrupole LC/MS/MS mass spectrometer (Applied Biosystems/Sciex Instruments) in positive electrospray ionization mode. The mass spectrometer was equipped with a Shimadzu LC-20AD liquid chromatograph (LC) and a SIL-20ACHT auto sampler. Chromatography was performed on a reverse-phase C18 column (Waters XBridge, 150 x 2.1 mm, 3 μm) with a mobile phase consisting of water/methanol (95:5, v/v) and 4mM dibutylamine acetate (eluent A), and water/acetonitrile (25:75, v/v) (eluent B). Multi-reaction monitoring (MRM) was carried out for the detection of the short-chain acyl-CoAs in standard solutions and biological samples. Acetyl-CoA was quantitated by comparison of the individual ion peak area with that of the internal standard. The analytical data were processed by Analyst Software (version 1.6.2).

##### ACC1 (radiometric)

Following pre-incubation in dephosphorylation treatment step, rACC1 was incubated at 37°C in a reaction mixture containing 20 mM HEPES (pH 7.1), 1 mM DTT, 1 mg/mL BSA, 5 mM NaHCO₃, 5 mM Mg-ATP (pH 7.1), 0.25 mM acetyl-CoA, and 0.225 µCi [¹⁴C]NaHCO₃ (American Radiolabeled Chemicals). At designated time points (0, 4, 8, 16, and 32 minutes), reactions were terminated by the addition of 8 N HCl. The mixtures were transferred to scintillation vials (Research Products International Corp.) and air-dried completely at 100°C. Dried samples were then dissolved in 1 mL of methanol, and [¹⁴C] radioactivity was quantified by liquid scintillation counting (Beckman Coulter) following the addition of scintillation cocktail (Research Products International Corp.).

##### FAS (radiometric)

Following pre-incubation in dephosphorylation treatment step, rFAS was incubated at 37°C in a reaction mixture containing 20 mM HEPES (pH 7.1), 1 mM DTT, 1 mg/mL BSA, 0.1 mM acetyl-CoA, 0.3 mM malonyl-CoA, 0.5 mM NADPH, and 0.02 µCi [¹⁴C]malonyl-CoA (American Radiolabeled Chemicals). At designated time points (0, 4, 8, 16, and 32 minutes), reactions were terminated by adding HCl/methanol (6:4, v/v) in 16 × 100 mm glass tubes (Corning). Ethanol and hexane were added at a volume ratio of 1:1:2 (HCl/methanol:ethanol:hexane), and the mixtures were vortexed thoroughly. Samples were centrifuged at 2,000 × g for 5 minutes at room temperature. The upper organic phase was transferred to scintillation vials and air-dried completely at RT. Dried lipids were dissolved in 1 mL of methanol, and [¹⁴C] radioactivity was quantified by liquid scintillation counting (Beckman Coulter) following the addition of scintillation cocktail (Research Products International Corp.).

##### ACL-ACC1-FAS (radiometric)

After pre-incubation in dephosphorylation treatment step, recombinant ACL (0.8 μg), ACC1 (0.4 μg), and FAS (3.2 μg) were incubated at 37 °C in a reaction mixture containing 20 mM HEPES (pH 7.1), 1 mM DTT, 1 mg ml⁻¹ BSA, 4 mM sodium citrate (pH 7.1), 5 mM NaHCO₃, 5 mM Mg-ATP (pH 7.1), 0.1 mM CoA, 1 mM NADPH, and 0.025 µCi [1,5-¹⁴C]citric acid (Revvity). At designated time points (0, 4, 8, 16, and 32 minutes), reactions were terminated by adding HCl/methanol (6:4, v/v) in 16 × 100 mm glass tubes (Corning). Ethanol and hexane were added at a volume ratio of 1:1:2 (HCl/methanol:ethanol:hexane), and the mixtures were vortexed thoroughly. Samples were centrifuged at 2,000 × g for 5 minutes at room temperature. The upper organic phase was transferred to scintillation vials and air-dried completely at RT. Dried lipids were dissolved in 1 mL of methanol, and [¹⁴C] radioactivity was quantified by liquid scintillation counting (Beckman Coulter) following the addition of scintillation cocktail (Research Products International Corp.).

##### ACSS2-ACC1-FAS (radiometric)

Following pre-incubation in dephosphorylation treatment step, recombinant ACSS2 (0.4 μg), ACC1 (0.4 μg), and FAS (3.2 μg) were incubated at 37 °C in a reaction mixture containing 20 mM HEPES (pH 7.1), 1 mM DTT, 1 mg ml⁻¹ BSA, 4 mM sodium acetate (pH 7.1), 5 mM NaHCO₃, 5 mM Mg-ATP (pH 7.1), 0.1 mM CoA, 1 mM NADPH, and 0.0225 μCi [1,2-¹⁴C]acetic acid (Revvity). At designated time points (0, 4, 8, 16, and 32 minutes), reactions were terminated by adding HCl/methanol (6:4, v/v) in 16 × 100 mm glass tubes (Corning). Ethanol and hexane were added at a volume ratio of 1:1:2 (HCl/methanol:ethanol:hexane), and the mixtures were vortexed thoroughly. Samples were centrifuged at 2,000 × g for 5 minutes at room temperature. The upper organic phase was transferred to scintillation vials and air-dried completely at RT. Dried lipids were dissolved in 1 mL of methanol, and [¹⁴C] radioactivity was quantified by liquid scintillation counting (Beckman Coulter) following the addition of scintillation cocktail (Research Products International Corp.).

##### Cytosol-based citrate-to-palmitate assay

Mouse livers collected after fasting-refeeding treatment were homogenized in Buffer I (20 mM HEPES, 250 mM sucrose, protease inhibitor cocktail (Millipore-Sigma), pH 7.1) at a liver-to-buffer ratio of 1:5 (w/v). Homogenates were centrifuged at 6,000 × g for 10 minutes at 4 °C. The resulting supernatant was subjected to further centrifugation at 180,000 × g for 60 minutes at 4 °C. The supernatant was then loaded on PD-10 desalting column (Cytiva) that pre-equilibrated with buffer J (20 mM HEPES, 50 mM KCl, 1 mM DTT, and protease inhibitor cocktail (Millipore-Sigma), pH 7.1) and flowed through by gravity. Liver cytosols were eluted in buffer J.

For pre-incubation, 300 μg of liver cytosol was incubated with 20 mM HEPES (pH 7.1), 1 mM MnCl₂, and phosphatase inhibitor cocktail at 30 °C for 15 minutes. Parallel samples without phosphatase inhibitor cocktail served as the dephosphorylated groups.

Following pre-incubation, liver cytosol samples were incubated at 37 °C for 15 minutes in a reaction mixture containing 20 mM HEPES (pH 7.1), 5 mM MgCl_2_, 1 mM DTT, 1 mg ml⁻¹ BSA, varying concentrations of sodium citrate (pH 7.1), 5 mM NaHCO₃, 5 mM ATP (disodium salt, pH 7.1), 0.1 mM CoA, 1 mM NADPH, and 0.05 μCi [1,5-¹⁴C]citric acid (Revvity). Reactions were terminated by adding 1 N HCl/methanol (6:4, v/v) in 16 × 100 mm glass tubes (Corning), followed by vortexing with ethanol and hexane. The final solvent ratio of quenching solution:ethanol:hexane was 1:1:2. Samples were centrifuged at 2,000 × g for 5 minutes at room temperature (RT), and the upper organic phase was collected, transferred to scintillation vials, and air dried completely at RT. Dried lipids were dissolved in 1 mL methanol, and [¹⁴C] radioactivity was quantified by liquid scintillation counting (Beckman Coulter) following the addition of scintillation cocktail (Research Products International Corp.).

##### TG synthesis in liver homogenates

The livers of fasted/refed rats were homogenized in buffer K (10 mM HEPES, 1 mM EDTA, 1 mM DTT, and protease inhibitor cocktail (Millipore-Sigma), pH 7.1) at a tissue-to-buffer ratio of 1:4 (w/v). The homogenate was filtered through gauze to remove unbroken tissue and then dialyzed against dialysis buffer (10 mM HEPES, 50 mM KCl, 1 mM DTT, 1 mM PMSF, pH 7.1) for 2 hours.

The liver homogenates were pre-incubated with various concentrations of MgCl_2_ (0, 2.5, 5, 10, 20 mM), 4 mM of Na-citrate buffer (pH 7.1), a phosphatase inhibitor cocktail, and 50 mM phosphate buffer (pH 7.1) for 5 minutes at 37°C. Parallel groups without phosphatase inhibitor cocktail and phosphate buffer served as dephosphorylated controls. After pre-incubation, samples were incubated for 30 minutes at 37 °C in a reaction mixture containing 1 mM DTT, 1 mg/mL BSA, 10 mM oleoyl-CoA, 100 mM glycerol 3-phosphate, and 0.2 μCi [¹⁴C]glycerol 3-phosphate (American Radiolabeled Chemicals). Reactions were terminated by adding 1 N HCl/methanol (6:4, v/v) in 16 × 100 mm glass tubes (Corning). The mixtures were then vortexed with phosphate-buffered saline (PBS) and Folch reagent (chloroform:methanol = 2:1, v/v). The final solvent ratio of HCl/methanol:PBS:Folch was 1:1:3. Samples were centrifuged at 2,000 × g for 5 minutes at room temperature (RT), and the lower organic phases were transferred to new glass tubes and air dried completely at RT. Dried lipids were reconstituted in 50 μL chloroform and vortexed. Extracted lipids, along with a TG marker, were loaded onto thin-layer chromatography (TLC) plates. Plates were developed in a TLC chamber saturated with a solvent system of hexane:diethyl ether:acetic acid (80:20:1, v/v/v). TG dots were visualized using iodine vapor, identified by comparison with the TG marker. TG dots were excised from the plates and transferred to scintillation vials. Scintillation cocktail (Research Products International Corp.) was added to each vial, followed by vortexing. [¹⁴C] radioactivity was quantified by liquid scintillation counting (Beckman Coulter).

##### Enzyme Synergy Assays (Recombinant ± Cytosol)

For dephosphorylation treatment, a mixture containing rACL (0.5 μg), rACC1 (0.25 μg), and rFAS (2 μg) or an equal volume of protein purification buffer (Buffer E) (used for the cytosol only groups) were incubated in a 25 μL total volume system with λ-phosphatase in the presence of 1 mM MnCl₂ at 30°C for 20 minutes. The resulting 22 μL reaction was combined with 20 μL of liver cytosol (300 μg) or cytosol preparation buffer (Buffer J) (used for the recombinant protein only groups) in a final volume of 50 μL and incubated at 30°C with 5 mM MgCl₂ for an additional 20 minutes.

To maintain proteins in a phosphorylated state, the same protein mixture or Buffer E was incubated under identical conditions with phosphatase inhibitor cocktail instead of λ-phosphatase. The resulting 22 μL reaction was then mixed with 20 μL of liver cytosol (300 μg) or Buffer J and incubated in a 50 μL system containing 5 mM MgCl₂, phosphatase inhibitor cocktail, and 50 mM phosphate buffer (pH 7.1) at 30°C for 20 minutes. Following the pre-incubation, samples were mixed with reaction buffer containing 20 mM HEPES (pH 7.1), 1 mM DTT, 1 mg/mL BSA, 4 mM sodium citrate (pH 7.1), 5 mM NaHCO₃, 5 mM Mg-ATP (pH 7.1), 0.1 mM CoA, 1 mM NADPH, and 0.025 μCi [1,5-¹⁴C]citric acid (Revvity) in a total volume of 75 μL. The reactions were incubated at 37°C for 15 minutes and terminated by adding 0.5 mL of cold methanol in 16 × 100 mm glass tubes (Corning). Next, 1.5 mL of methyl tert-butyl ether (MTBE) was added, and the samples were vortexed thoroughly and left to stand at RT for 5 minutes. Then, 0.5 mL of ddH₂O was added, followed by another round of vortexing and a 5-minute incubation at RT. Phase separation was achieved by centrifugation at 2,000 × g for 5 minutes at RT. The upper organic phase was transferred to scintillation vials and air-dried completely at RT. Dried lipids were dissolved in 1 mL of methanol, and [¹⁴C] radioactivity was quantified by liquid scintillation counting (Beckman Coulter) after the addition of scintillation cocktail (Research Products International Corp.).

##### Phosphatase inhibitor cocktail preparation

The phosphatase inhibitor cocktail was prepared fresh using the following recipe: sodium fluoride (200 mM), imidazole (200 mM), sodium molybdate (115 mM), sodium orthovanadate (200 mM), sodium tartrate dihydrate (400 mM), sodium pyrophosphate (100 mM), and β-glycerophosphate (100 mM) in ddH₂O (final volume 20 mL). Sodium orthovanadate was prepared separately by dissolving 1.84 g of powder in 50 mL ddH₂O, adjusting the pH to 10, briefly boiling in a microwave (10 sec), cooling on ice to room temperature, and repeating the cycle three times. The solution was adjusted to 50 mL, aliquoted, and stored at –20°C until use.

##### Quantification and statistical analysis

Sample sizes (n) are indicated in the figure legends. Technical replicates are also noted in the figure legends when applicable. Unless otherwise stated, data are presented as mean ± SEM. Statistical analyses were performed using GraphPad Prism 10. The statistical test and model used for each experiment are specified in the corresponding figure legend. A significance threshold of α = 0.05 was used.

For co-localization analyses, images were quantified using Fiji/ImageJ (Coloc2). Pearson’s correlation coefficients were reported as values above the Costes threshold. Thresholded Manders’ coefficients (tM1 and tM2), Costes significance testing, and 2D intensity histograms were generated using Coloc2 and are provided in the Supplemental Information.

#### Supplemental item titles

**Figure S1. Representative regions of interest (ROIs) and pixel-intensity scatter plots used for colocalization analysis of ACC1, S14, and MIG12, related to Figure 1**.

**Figure S2. Sequential formation of the ACC1-containing lipogenic complex around LDs, related to Figure 2**.

**Figure S3. Electron microscope images, related to Figure 3**.

**Figure S4. Phosphorylated ACC levels in liver lysates from rats refed the FFD, related to** Figure 4.

**Figure S5. Physical association of ACC1 and ACC2 and regulation of TG synthesis by phosphorylation in an MgCl_2_-dependent manner, related to Figure 5**.

**Figure S6. Lipogenic metabolon formation facilitates FA production, related to Figure 6**.

**Figure S7. DAVID functional annotation of ACC1-dependent interactors from ACC1, MIG12, and S14 immunoprecipitates, related to Figure 7**.

**Supplemental Table 1. Pearson’s and Manders’ colocalization coefficients together with Costes significance statistics for ACC1–MIG12, S14–MIG12, and S14–ACC1, related to Figure 1**.

**Supplemental Table 2. Quantitative co-localization analysis of ACC1 with cytosolic lipogenic enzymes in primary hepatocytes, related to Figure 2**.

**Supplemental Table 3. Original data of MS analysis of ACC1-dependent interactors from ACC1, MIG12, and S14 immunoprecipitates, related to Figure 7**.

