## Supplemental Figures and Figure Legends for "Identification and Regulation of a Hepatic Lipogenic Metabolon"

### Figure S1

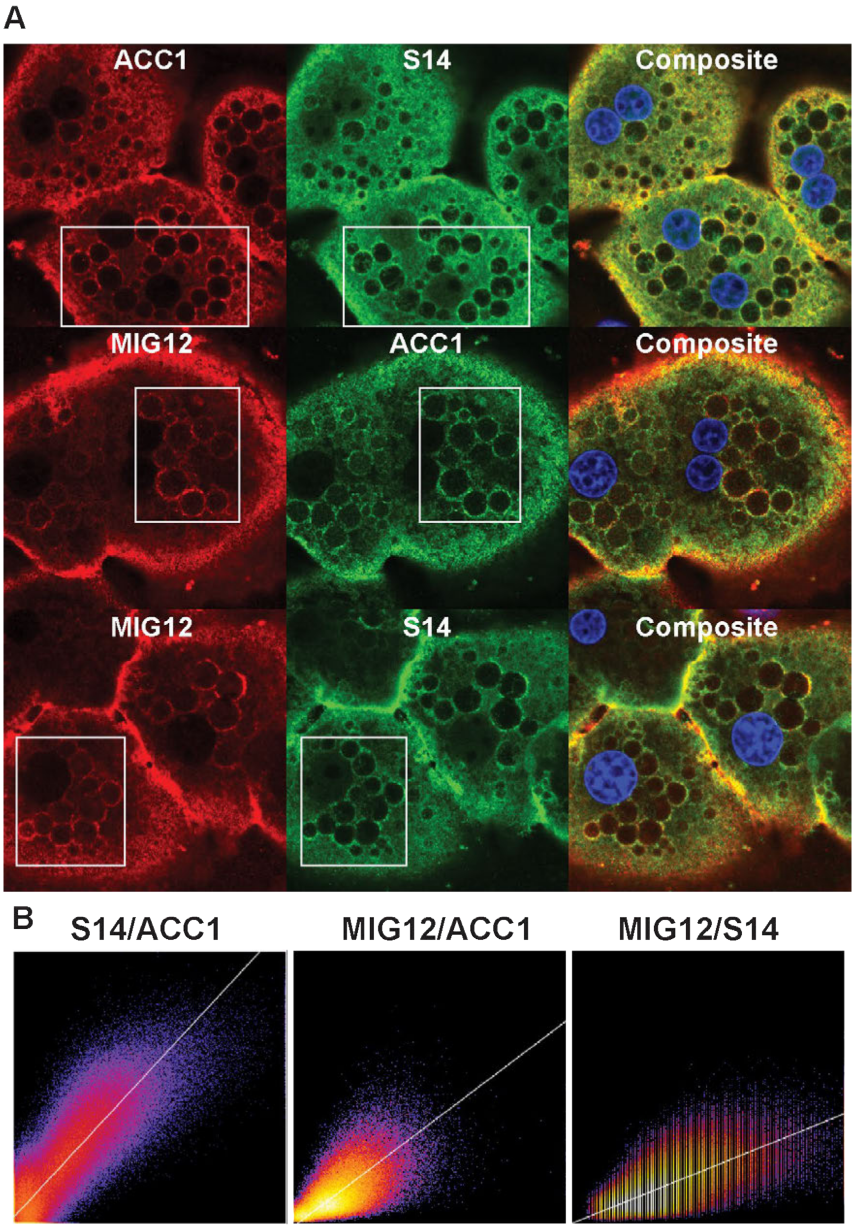


**Figure S1. Representative regions of interest (ROIs) and pixel-intensity scatter plots used for colocalization analysis of ACC1, S14, and MIG12, related to Figure 1.**

**(A)** Primary hepatocytes isolated from rats refed a fat-free, high-carbohydrate diet (FFD) were stained for ACC1, MIG12, and S14 and imaged by confocal microscopy. For each enzyme pair, colocalization was quantified within lipid droplet–enriched regions, where lipogenic enzymes preferentially accumulate. White boxes indicate the ROIs used for analysis.

**(B)** Bottom panels show 2D pixel-intensity histograms (Coloc2, Fiji) for each enzyme pair, illustrating the correlation between fluorescence intensities of the two channels. Diagonal clustering of pixels indicates positive spatial correlation. Pearson's correlation coefficients (above Costes threshold) for each pair are reported in Figure 1B, while full colocalization metrics—including non-thresholded and thresholded Pearson's R, Manders coefficients, and Costes significance testing—are provided in Supplementary Table 1.

#### Figure S2

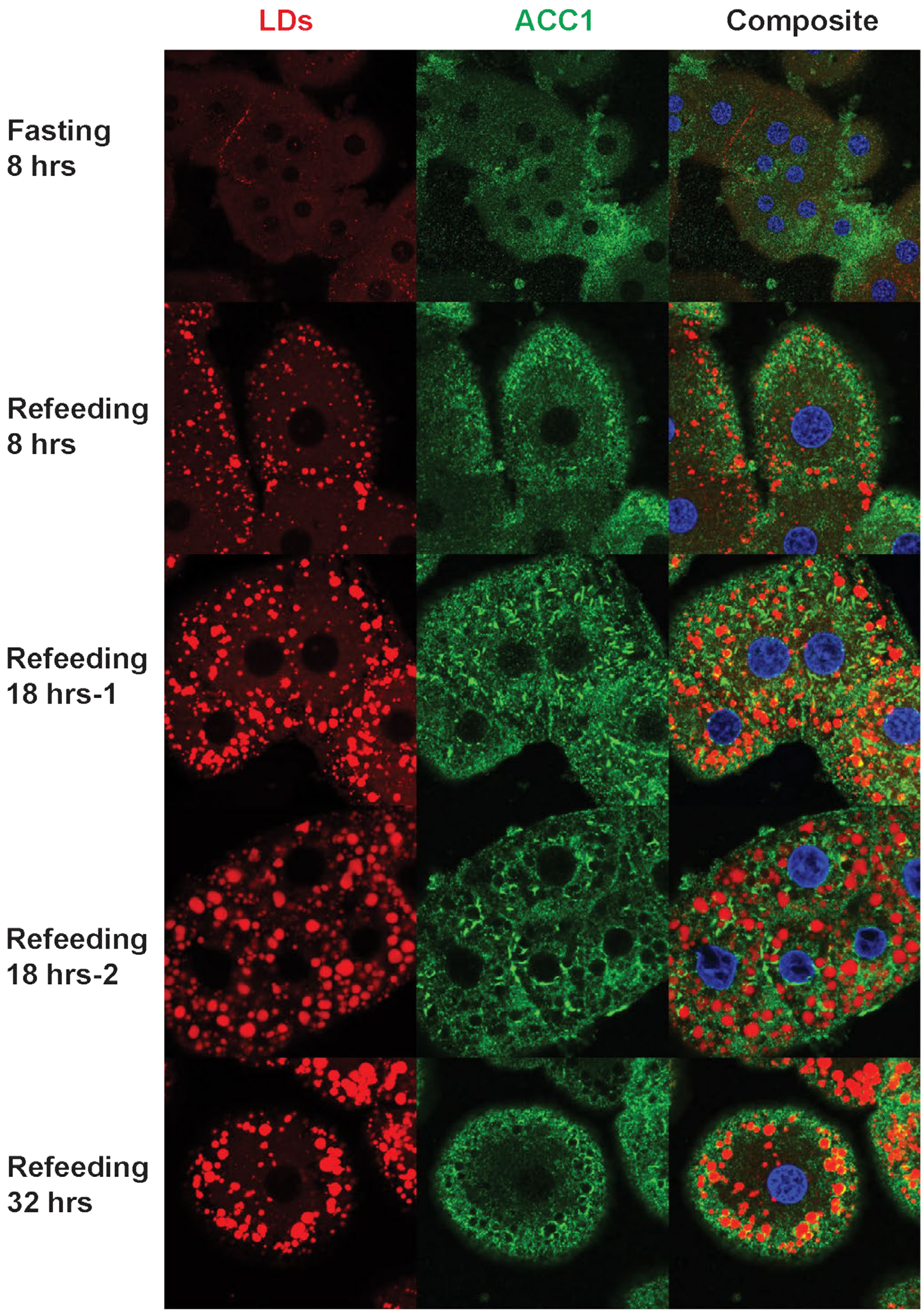


**Figure S2. Sequential formation of the ACC1-containing lipogenic complex around LDs, related to Figure 2.**

Primary hepatocytes were prepared from chow-fed rats and cultured in low glucose (5 mM)/low insulin (1 nM) medium (fasting) or high glucose (25 mM)/high insulin (100 nM) medium (refeeding). Immunofluorescence images were obtained at indicated times, and antibodies were labeled as described in STAR Methods.

#### Figure S3

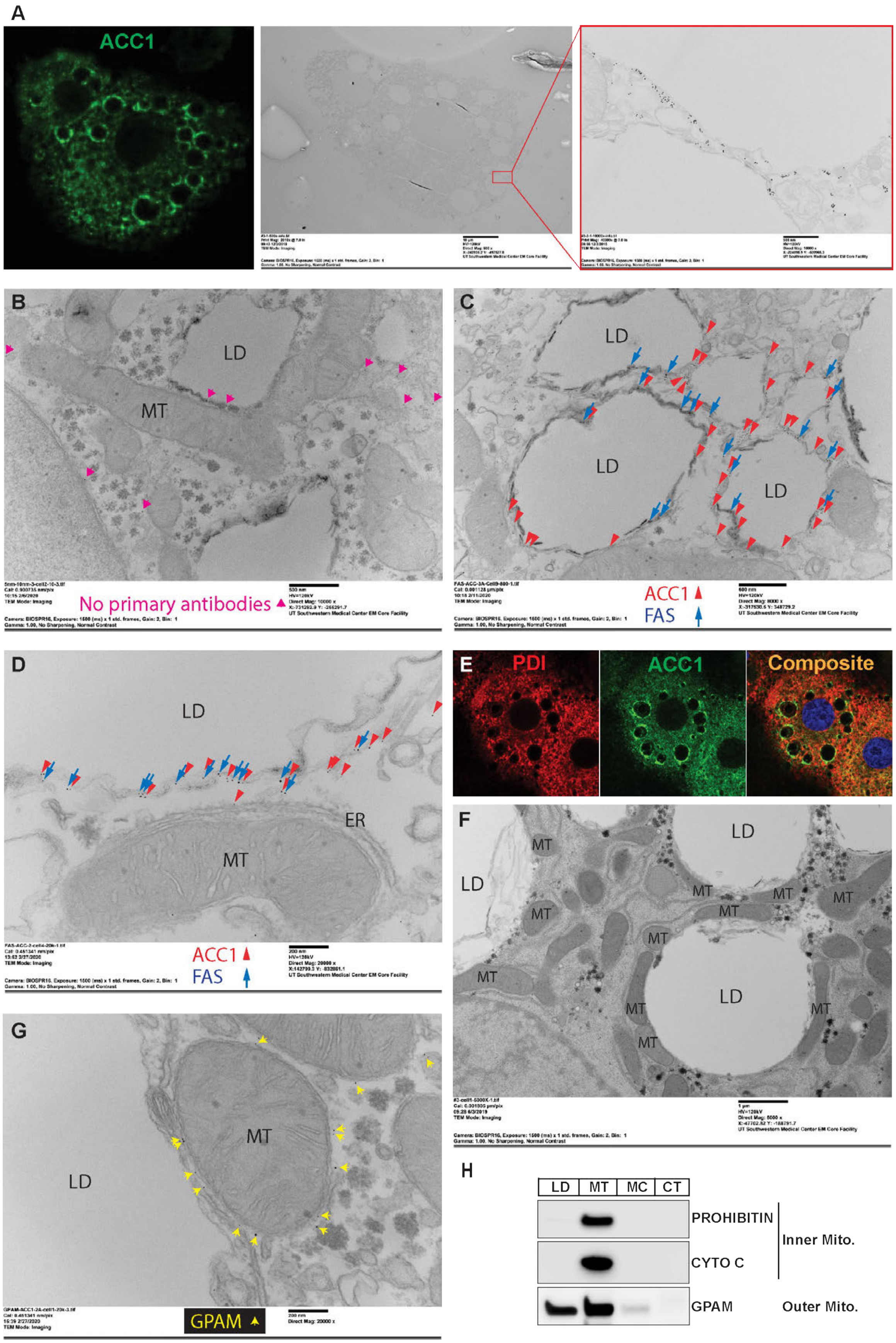
A
ACC1
B
LD
MT
No primary antibodies
C
LD
LD
LD
ACC1
FAS
D
LD
ER
MT
ACC1
FAS
E
PDI
ACC1
Composite
F
LD
LD
LD
MT
G
MT
LD
GPAM
H
LD
MT
MC
CT
PROHIBITIN
CYTO C
Inner Mito.
GPAM
Outer Mito.

**Figure S3. Electron microscope images, related to Figure 3.**

**(A)** Immunofluorescence and electron microscope images were obtained from the same primary hepatocytes from rats refed the FFD to demonstrate the localization of ACC1 around LDs.

**(B-D)** Immunogold-EM images of primary hepatocytes from rats refed the FFD. (B) No primary antibodies (secondary antibody only) to assess nonspecific binding. (C–D) Co-localization of ACC1 (5 nm gold) and FAS (10 nm gold) in two representative regions, including an LD-rich area (C) and an area where mitochondria and ER are in close proximity to LDs (D).

**(E)** Immunofluorescence was taken from the primary hepatocytes from rats refed with the FFD to demonstrate the localization of PDI and ACC1.

**(F)** Numerous mitochondria were observed in close proximity to LDs in primary hepatocytes from rats refed the FFD.

**(G)** Localization of GPAM on mitochondria and the interface between mitochondria and lipid droplets in primary hepatocytes from rats refed the FFD.

**(H)** Immunoblot analysis following subcellular fractionation of the specified inner and outer mitochondrial proteins in liver lysates from rats refed the FFD.

### Figure S4

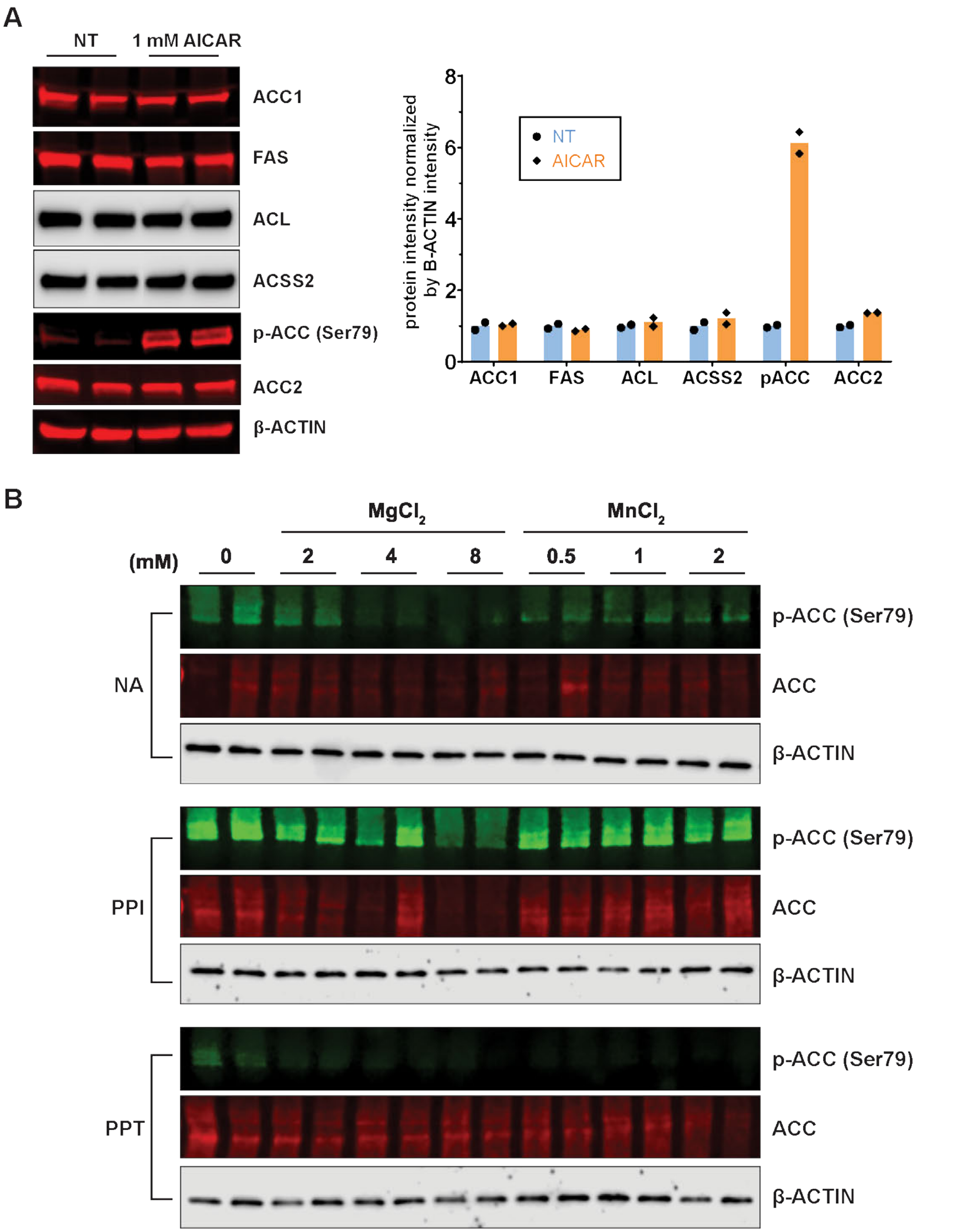


**Figure S4. Phosphorylated ACC levels in liver lysates from rats refed the FFD, related to Figure 4.**

**(A)** Cell lysates were prepared from primary rat hepatocytes incubated with 100 nM insulin or 1 mM AICAR for 30 minutes. Equal aliquots of cell lysate proteins were subjected to SDS-PAGE for immunoblot analysis.

**(B)** Liver lysates from rats refed with the FFD were incubated with the phosphatase inhibitor (PPI) cocktail or λ-phosphatase (PPT) under various $MgCl_2$ (0, 2, 4, 8 mM) or $MnCl_2$ (0, 0.5, 1, 2 mM) concentrations at 30°C for 30 min. Groups without either PPI or PPT served as controls (NA). Equal amounts of liver lysate proteins were subjected to SDS-PAGE for immunoblot analysis. ACC immunoblots were performed using a fluorescently labeled streptavidin conjugate.

#### Figure S5

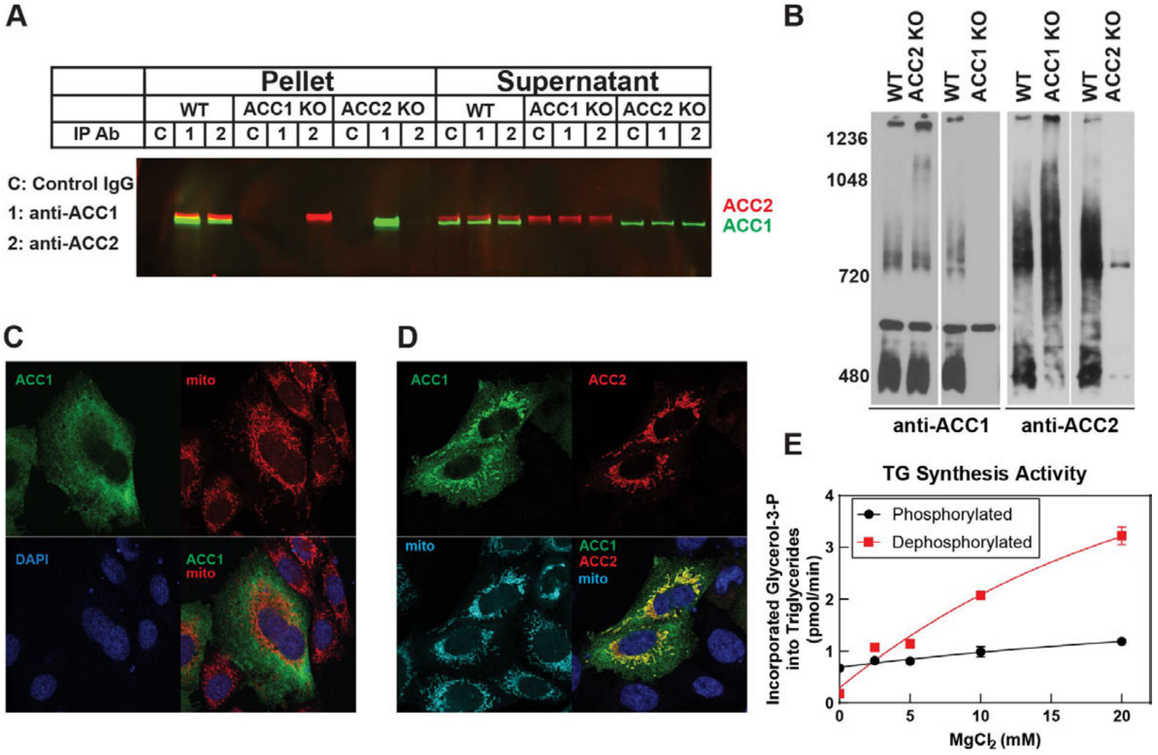


**Figure S5. Physical association of ACC1 and ACC2 and regulation of TG synthesis by phosphorylation in an $MgCl_2$-dependent manner, related to Figure 5.**

**(A)** ACC1 or ACC2 were immunoprecipitated using an anti-ACC1 or anti-ACC2 antibody from the liver lysates prepared from WT, ACC1 hepatocyte KO, or ACC2 hepatocyte KO mouse livers. Pellets and supernatants following the IP were loaded on a 4-15% SDS-PAGE gel and immunoblot analysis was carried out using the anti-ACC1 and ACC2 antibodies.

**(B)** BN-PAGE was performed essentially as described in Kim et al. [1] Briefly, cytosolic fractions from WT, ACC1 LKO, and ACC2 LKO livers were pooled within each genotype, and 30 µg of protein was resolved on a 3.5–8% BN-PAGE gel. Following transfer to PVDF membranes, blots were probed with the indicated anti-ACC1 or anti-ACC2 antibodies.

**(C-D)** CHO-K1 cells were transfected with ACC1-3x HA (5 µg) and ACC2-5x Myc (2 µg) alone or in combination and cultured for 24 hours at 37 °C and 8.8% CO2. Prior to fixation, cells were treated with 300 nM MitoTracker for 30 minutes. Cells were fixed with 4% PFA/PBS, quenched in 10% glycine, and permeabilized with 0.2% IGEPAL. After blocking at room temperature, cells were stained with antibodies against HA (ACC1) and Myc

(ACC2) for 2 hours. Secondary antibodies were incubated with cells for 1 hour, and coverslips were mounted in DAPI mounting media overnight.

**(E)** Liver lysates from rats refed with the FFD were pre-incubated with various concentrations of $MgCl_2$ for 5 min at 37°C in the presence or absence of 50 mM Na-phosphate buffer (pH 7.1), which served as a source of inorganic phosphate to inhibit phosphatase activity. Afterward, the lysates were incubated with 1 mM DTT, 1 mg/ml BSA, 10 mM oleoyl-CoA, 100 mM glycerol-3-phosphate, and 18 µM [$^{14}$C]glycerol 3-phosphate for 30 min at 37°C. Lipids were extracted, and TGs were isolated via thin-layer chromatography (TLC). The incorporation of $^{14}$C-glycerol-3-P was used to trace newly synthesized TG. The TG synthesis activities were measured and plotted as a function of $MgCl_2$ concentration. Data points represent individual measurements from duplicate experiments. Detailed experimental protocols are provided in STAR Methods.

### Figure S6

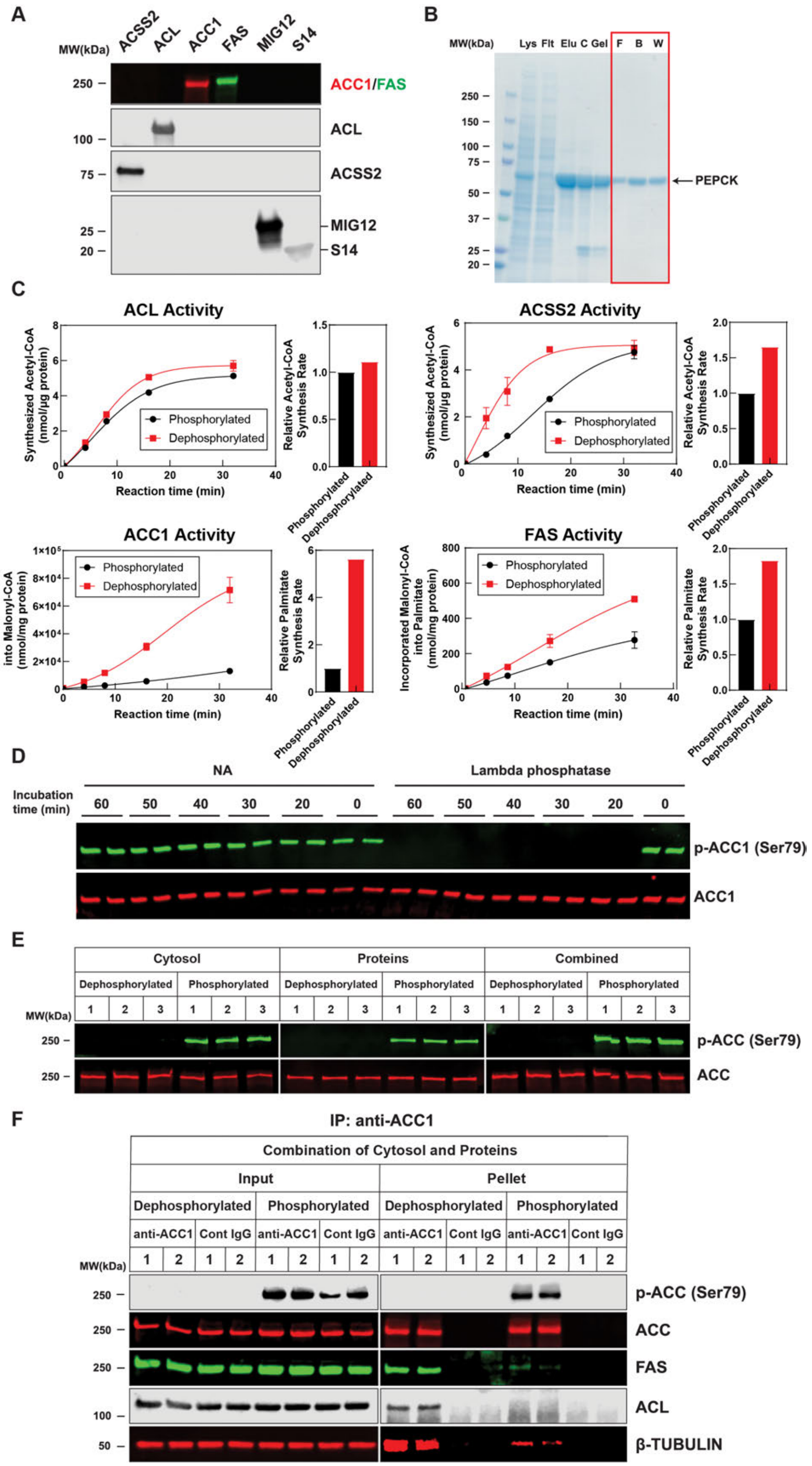
A
MW(kDa)
ACSS2
ACL
ACC1
FAS
MIG12
S14
250
100
75
25
20
ACC1/FAS
ACL
ACSS2
MIG12
S14
B
MW(kDa)
Lys
Flt
Elu
C
Gel
F
B
W
250
150
100
75
50
37
25
20
PEPCK
C
ACL Activity
Synthesized Acetyl-CoA (nmol/µg protein)
Reaction time (min)
Phosphorylated
Dephosphorylated
Relative Acetyl-CoA Synthesis Rate
ACSS2 Activity
ACC1 Activity
Incorporated HCO3- into Malonyl-CoA (nmol/mg protein)
Relative Palmitate Synthesis Rate
FAS Activity
Incorporated Malonyl-CoA into Palmitate (nmol/mg protein)
D
NA
Lambda phosphatase
Incubation time (min)
60
50
40
30
20
0
p-ACC1 (Ser79)
ACC1
E
Cytosol
Proteins
Combined
Dephosphorylated
Phosphorylated
p-ACC (Ser79)
ACC
F
IP: anti-ACC1
Combination of Cytosol and Proteins
Input
Pellet
anti-ACC1
Cont IgG
p-ACC (Ser79)
ACC
FAS
ACL
β-TUBULIN

**Figure S6. Lipogenic metabolon formation facilitates FA production, related to Figure 6.**

**(A)** Immunoblot analysis of purified proteins was performed using antibodies specific to each protein, respectively. ACC1 immunoblots were performed using a fluorescently labeled streptavidin conjugate.

**(B)** Proteins from different fractions produced during the purification of recombinant PEPCK were subjected to SDS-PAGE for Coomassie blue staining. The final recombinant PEPCK was obtained from the combination of the fractions framed in red. Lys = Cell lysates from HEK293 cells; Flt = Flow through from Ni-NTA resin; Elu = Elution fractions from the Ni-NTA resin (500 mM imidazole); C = Fraction after TEV cleavage; Gel = Gel filtration elution; F = Flow through of gel filtration elution from Ni-NTA resin; B = Flow through of binding buffer from Ni-NTA resin (5 mM imidazole); W = Flow through of wash buffer from Ni-NTA resin (50 mM imidazole).

**(C)** Enzymatic activity measurements of the indicated recombinant proteins were performed as described in STAR Methods. Data points represent individual measurements from duplicate experiments. Relative synthesis rates were measured based on the Vmax of each enzymatic reaction.

**(D)** Recombinant ACC1 was incubated with λ-phosphatase and 1 mM $MnCl_2$ for 0, 20, 30, 40, 50, and 60 min and subjected to SDS-PAGE for immunoblot analysis. NA indicates rACC1 incubated with 1 mM $MnCl_2$ for 0, 20, 30, 40, 50, and 60 min. ACC1 immunoblots were performed using a fluorescently labeled streptavidin conjugate.

**(E)** Equal amounts of samples of liver cytosol (from rats refed with the FFD) alone group, recombinant proteins (rACL, rACC1, and rFAS) alone group, and combined liver cytosol–recombinant protein group from Fig. 6D and 6E after pre-treatment were subjected to SDS-PAGE for immunoblot analysis. ACC immunoblots were performed using a fluorescently labeled streptavidin conjugate.

**(F**) A combination of liver cytosol (from rats refed with the FFD) and recombinant proteins (rACL, rACC1, and rFAS) were pre-treated under dephosphorylated and phosphorylated conditions following the same protocol described in Figures 6D and 6E. The ACC1 antibody was incubated with the mixture for 1 hour at RT and ACC1 was immunoprecipitated. Control IgG was used for the control IP. The immunoprecipitation protocol is described in STAR Methods. Proteins were subjected to SDS-PAGE for immunoblot analysis. ACC immunoblots were performed using a fluorescently labeled streptavidin conjugate.

#### Figure S7

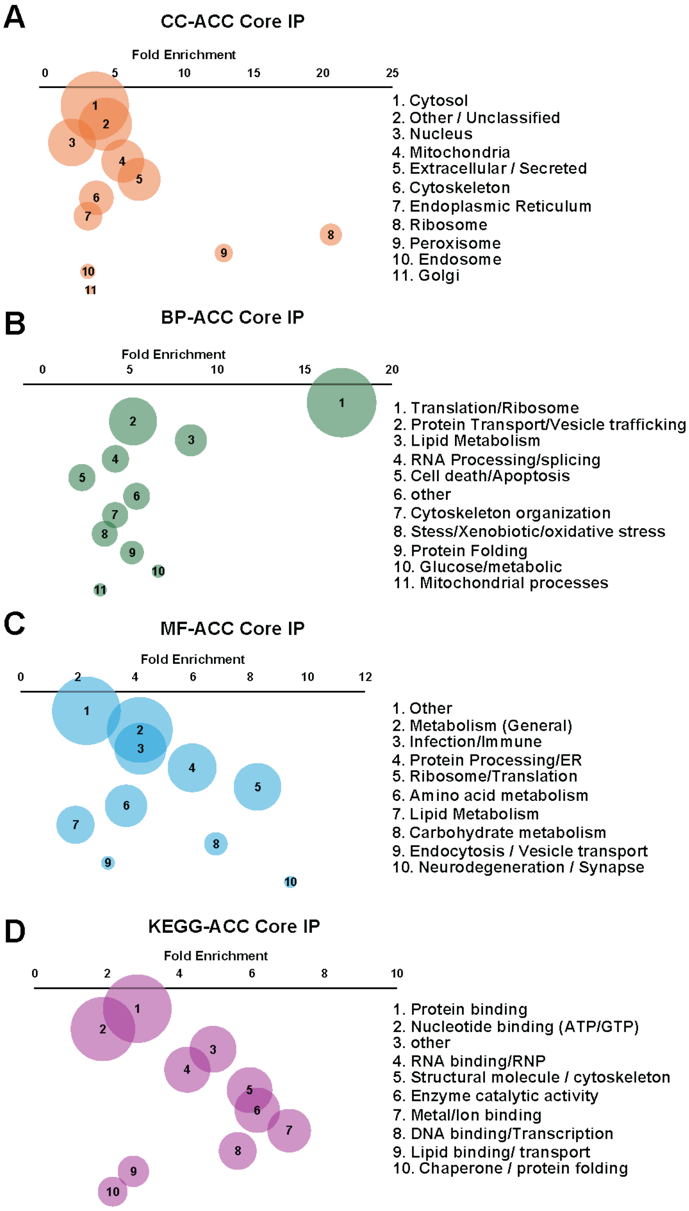


**Figure S7. DAVID functional annotation of ACC1-dependent interactors from ACC1, MIG12, and S14 immunoprecipitates, related to Figure 7.**

Proteins co-immunoprecipitated with ACC1, MIG12, and S14 from WT and ACC-deficient (ACC-KO) liver lysates were quantified by mass spectrometry, and proteins reduced by >20-fold in ACC-KO were defined as ACC1-dependent interactors. Gene symbols from this filtered list were uploaded to DAVID 6.8 (https://david.ncifcrf.gov/; Mus musculus background), and enrichment outputs from GO_CC, GO_BP, GO_MF, and KEGG modules were exported and manually consolidated into broader functional categories. Bubble plots display fold enrichment on the x-axis and gene count as bubble size.

**(A)** GO_CC analysis showing enriched subcellular compartments; numeric labels correspond to merged category names (right).

**(B)** GO_BP analysis highlighting dominant biological processes grouped into higher-order functional terms.

1. Kim, C.W., Moon, Y.A., Park, S.W., Cheng, D., Kwon, H.J., and Horton, J.D. (2010). Induced polymerization of mammalian acetyl-CoA carboxylase by MIG12 provides a tertiary level of regulation of fatty acid synthesis. Proc Natl Acad Sci U S A *107*, 9626-9631. 10.1073/pnas.1001292107.
